# The use of high throughput phenotyping for assessment of heat stress-induced changes in Arabidopsis

**DOI:** 10.1101/838102

**Authors:** Ge Gao, Mark A. Tester, Magdalena M. Julkowska

## Abstract

The worldwide rise in heatwave frequency poses a threat to plant survival and productivity. Determining the new marker phenotypes that show reproducible response to heat stress and contribute to heat stress tolerance is becoming a priority. In this study, we describe a protocol focusing on the daily changes in plant morphology and photosynthetic performance after exposure to heat stress using an automated non-invasive phenotyping system. Heat stress exposure resulted in an acute reduction of quantum yield of photosystem II and increased leaf angle. In the longer term, exposure to heat also affected plant growth and morphology. By tracking the recovery period of WT and mutants impaired in thermotolerance (hsp101), we observed that the difference in maximum quantum yield, quenching, rosette size, and morphology. By examining the correlation across the traits throughout time, we observed that early changes in photochemical quenching corresponded with the rosette size at later stages, which suggests the contribution of quenching to overall heat tolerance. We also determined that 6h of heat stress provides the most informative insight in plant responses to heat, as it shows a clear separation between treated and non-treated plants as well as WT and hsp101. Our work streamlines future discoveries by providing an experimental protocol, data analysis pipeline and new phenotypes that could be used as targets in thermotolerance screenings.

## 1. Introduction

Globally, the last decade was the warmest since the 19th century, and resulted in record-breaking heat waves in many parts of the world (Coumou et al., 2013; Schiermeier, 2019). Heat stress leads to a reduction of plant performance and productivity at all developmental stages, making the heatwaves a serious threat to agriculture. However, the majority of the efforts in heat stress research focus either on early seedling development, scoring survival or hypocotyl elongation (Larkindale et al., 2005; Silva-Correia et al., 2014; McLoughlin et al., 2016) or reproductive stages (Zhou et al., 2017; Fan et al., 2018), where the pollen viability is reduced by high temperatures. The handful of studies focusing on the heat stress responses at the vegetative development stage (Rodríguez et al., 2015; Xu et al., 2017; Cheabu et al., 2018) show that heat tolerance at vegetative stage contributes to resilience at the reproductive stage. Therefore, understanding the changes caused by heat stress and breeding for heat tolerance at all developmental stages is essential to ensure future sustainable food supply.

*Arabidopsis thaliana* has been widely used in screenings for thermotolerance, predominately focusing on seedling viability (Li et al., 2007; Gao et al., 2008; Suzuki et al., 2008), hypocotyl elongation (Hong and Vierling, 2000; Charng et al., 2007), or seed germination (Silva-Correia et al., 2014) on agar plates. As heat tolerance relies on multiple processes, quantification of simple traits, determined by the ease of phenotyping rather than physiological importance, does not provide the best tools capturing the complexity of the responses, e.g. plant cooling capacity, growth recovery, and maintenance of photosynthesis, which all contribute to the diversity of thermotolerance mechanisms. Continuous monitoring of plant growth after heat exposure via non-destructive methods, such as RGB, thermal imaging and chlorophyll fluorescence, provide insight into physiological responses corresponding to photosynthetic efficiency and plant cooling abilities which cannot be scored by eye. The automated and environmentally controlled system enables time-efficient screening of large populations in a single experiment. Phenotypic traits, such as plant size, temperature, and photosynthetic efficiency have been successfully applied to evaluate plant performance under drought (Jansen et al., 2009; Chen et al., 2014), salinity (Awlia et al., 2016), and chilling (Jansen et al., 2009) stress, but to our knowledge, no such study has been conducted on study heat stress response.

HSP101, a molecular chaperone involved in protein disaggregation was one of the earliest genes identified to have a crucial role in thermotolerance in Arabidopsis, with no detrimental effects on normal growth or development in the absence of stress (Queitsch et al., 2000; Hong and Vierling, 2001). Homologs of HSP101 were identified and characterized for their role in heat stress response in maize, soybean, wheat, tobacco and pea, kidney bean (Keeler et al., 2000; Katiyar-Agarwal et al., 2001). As such, *hsp101* mutant showed a severe reduction in heat tolerance compared to wild-type in terms of survival, however the broader knowledge about the processes compromised in this mutant during the heat exposure are unknown.

In this study, we investigate the feasibility of applying RGB, kinetic chlorophyll fluorescence and infrared imaging for evaluating heat stress response in Arabidopsis. We developed a physiologically relevant heat-imposition protocol for the vegetative stage of Arabidopsis plants based on the significant changes observed for multiple traits. Additionally, by studying the heat-induced changes in WT and *hsp101* mutant, we were able to identify additional traits that might indicate compromised heat stress tolerance. By applying machine learning, we identified that maintenance of photochemical quenching immediately after stress application could be potentially used as an indicator for heat stress tolerance, as it corresponded with the increase in plant size at the later time points. This work provides a primer for future studies using high-throughput phenotyping platforms, uncovering novel components of heat stress tolerance.

## 2. Materials and Methods

### 2.1 Plant materials and growth conditions

Seeds of Arabidopsis wildtype Col-0 (CS60000) and *hsp101* (AT1G74310; *hot 1-3*, NASC ID: N16284) were stratified for three days at 4°C in the dark and germinated in controlled environment in PSI growth room (Photon Systems Instruments, Czech Republic). The environmental setting of PSI growth room) was set at 22°C (sensor sensitivity range: ± 0.1°C), with a relative humidity of 60% (sensor sensitivity range: ± 1%) and 400 ppm (sensor sensitivity range: ±100 ppm) of CO_2_. At four-leaf stage (day 14 after sowing), healthy seedlings with similar size were transferred into PSI standard pots (6 cm × 6 cm × 9.5 cm) filled with 100 g (± 1.0 g) of the growing mix (SunGro Horticulture Metro-Mix 360, MA, USA), placed into PSI trays (5 × 4 pots per tray) and registered into the PlantScreen™ system. All pots were automatically weighed and watered every day to reach and maintain the weight of 130 g. Plants were grown under cool-white LED panel with a 16 h/8 h light/dark cycle, the light intensity received at plant rosette level is ∼150 μmol m^−2^ s^−1^. All plants were kept in the PSI growth room during the experiment except during the heat treatment.

### 2.2 Heat stress treatment and phenotyping experimental design

At day 22 after sowing, we subjected plants to 3 h, 6 h or 9 h of heat stress and control treatments (**Figure 1**). For each treatment group (3 h, 6 h, 9 h or control), there are two trays each containing 10 wildtype Col-0 and 10 *hsp101* plants next to each other in an evenly distributed design. Heat stress was applied by placing the 9 h, 6 h and 3 h treatment trays into a pre-heated Percival growth chamber (Model CU36-L5, Percival Scientific, IA, USA) with white lights on (45°C, ∼120 μmol m^−2^ s^−1^) from 9 am to 6 pm, from 12 pm to 6 pm, and 3 pm to 6 pm respectively. Two trays used as “control treatment” with the same composition of genotypes remained at the PSI growth room. After heat stress application, six trays were transferred back to the PSI growth room. The transfer of the trays between the phenotyping facility and the heated growth chamber lasted between 2-3 minutes. Plants were imaged using chlorophyll fluorescence, RGB and infrared cameras daily at 7 pm starting from the day before the heat stress application (DAS-1) until one week after (**Figure 1**). Using image-based analysis, we acquired a variety of traits reflecting plant growth, photosynthetic efficiency, rosette morphology, and temperature in Col-0 and *hsp101* plants and explored these phenotypes for phenotypic plasticity in response to heat stress and their feasibility for large thermotolerance screening. In total, we screened 160 individual plants, with 20 biological replicates per genotype per treatment, to develop an understanding of changes induced by treatment, genotype and interaction between them for various traits.

**Figure 1.**
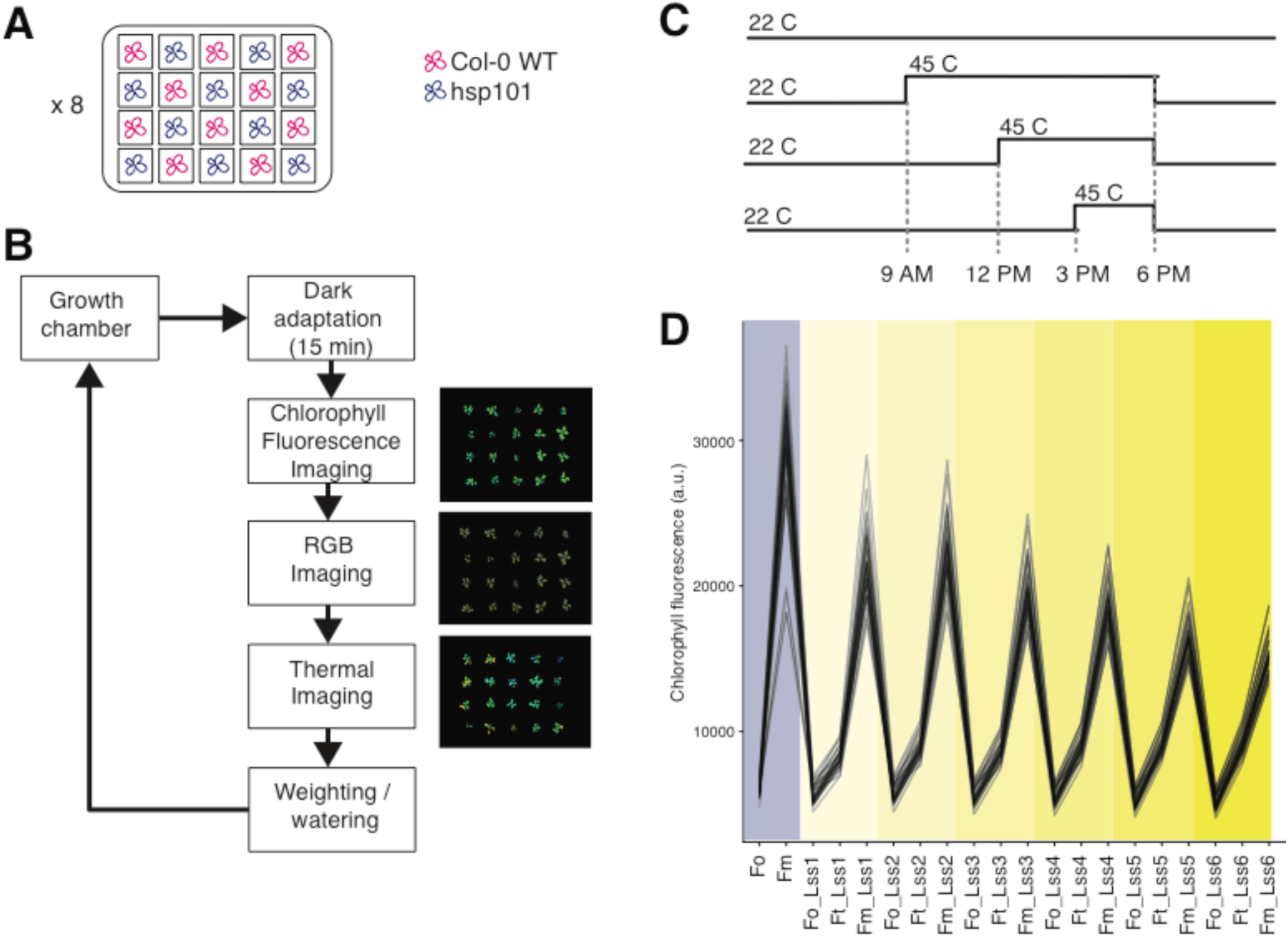
Schematic of the experimental setup. **(A)** Col-0 (WT) and *hsp101* seedlings were grown in standardized pots supplied by the PlantScreen™ phenotyping system using a checker-board design across 8 trays, with 10 Col-0 and *hsp101* seedlings in each tray. The environment in the growth room was set to a 16/8 h day/night cycle, with 22°C and 60 % relative humidity. **(B)** The phenotyping protocol. Each tray underwent an initial 15 min dark-adaptation period inside the adaptation chamber, followed by chlorophyll fluorescence, red green blue (RGB), and thermal imaging, with automatic weighing and watering before returning to the growth chamber. **(C)** The heat stress imposition protocol. 22 days after sowing, two trays of plants were kept in the growth chamber as control and the other six trays were moved into the pre-heated 45°C Percival chamber at 9 am, 12 pm and 3 pm for the 9 h, 6 h, and 3 h heat treatment respectively. All treated six trays were returned to the growth chamber at 6 pm and imaged daily at 7 pm starting from the day before the heat stress application (DAS-1) until one week after. **(D)** The overview of the chlorophyll fluorescence protocol executed using the dark-adapted plants. The minimal (Fo) and maximal (Fm) fluorescence are measured directly after dark adaptation, followed by gradual exposure to increasing light intensities of 95, 210, 320, 440, 555 and 670 µ mol m^−2^ s^−1^, corresponding to Lss 1, 2, 3, 4, 5, and 6 respectively, where the minimal (Fo’) and steady-state fluorescence are determined. At each light intensity plants are exposed to a saturating light flash, which allows measuring the maximum fluorescence at light-adapted state for given intensity (Fm’).

### 2.3 Imaging-based phenotypic measurements

Plant imaging was initiated one day before the heat application (DAS-1) to provide a baseline for the analysis and was performed daily at 7 pm until 7 days after the heat application (DAS 7). Each imaging round consisted of an initial 15 min dark-adaptation period inside the acclimation channel, followed by chlorophyll fluorescence, red green blue (RGB) colored and infrared (IR) imaging. For each imaging round, the phenotyping time for all trays was about 80 mins. Lighting conditions, plant positioning, and camera settings were fixed throughout the experiment.

Chlorophyll fluorescence imaging unit in PlantScreen™ Systems constructed by Photon System Instruments (PSI, The Czech Republic) measures the re-emitted light approximating the photosynthetic performance of plants’ photosystem II. The light curve protocol from PSI was applied to provide detailed information on fluorescence kinetics during the heat stress recovery (Henley, 1993; Rascher *et al.*, 2000), a detailed protocol can be found in **Figure S1**. After 15 minutes of dark adaptation, the initial flash of light was applied to measure the minimum fluorescence (F_o_), followed by a saturation pulse to determine the maximum fluorescence (F_m_) in the dark-adapted state. Next, six 60 s intervals of cool-white actinic light with increasing intensity of 95, 210, 320, 440, 555 and 670 µ mol m^−2^ s^−1^ was applied to record the chlorophyll fluorescence signal at the end of each actinic light period as the steady-state fluorescence in light-adapted state (F_t_’), followed by signal measured at the saturation pulse as the maximal fluorescence in the light-adapted state (F_m_’). Based on those basic chlorophyll fluorescence signals, variable fluorescence during dark-adapted state (F_v_, calculated as F_m_-F_o_), the maximum quantum yield of PSII photochemistry (QY max calculated as F_v_/F_m_), variable fluorescence in steady-state (F_v_’, calculated as F_m_’ – F_o_’), steady-state PSII quantum yield in the light-adapted state (QY, calculated as (F_m_’ – F_t_)/F_m_’), steady-state non-photochemical quenching (NPQ, calculated as (F_m_ – F_m_’)/F_m_’), coefficient of non-photochemical quenching during light-adapted state (qN, calculated as (F_m_-F_m_’)/(F_m_-F_o_’)), coefficient of photochemical quenching during light-adapted state (qP, calculated as (F_m_’-F_t_)/(F_m_’-F_o_’)) (**Table S1**).

Plant growth and morphological traits were obtained by RGB imaging unit using a 5-megapixel RGB camera (SV-0814H, VS Technology) mounted above the passing trays, providing the top-view image. The color images were processed in PlantScreen™ Analyser software to develop plant masks, used for chlorophyll fluorescence imaging and extraction of morphological parameters such as rosette compactness, eccentricity, and roundness, calculated by the PlantScreen™ Analyser software. Thermal camera (A615, FLIR) was used to detected infrared radiation, which is converted into an electronic signal and subsequently processed to provide thermal images. The RGB mask was used to extract pixels belonging to individual plants on the thermal image, and output the average surface temperature of the rosette area.

### 2.4 Data Analysis

Raw data were retrieved from the PlantScreen™ Analyser software, data visualization performed in R (http://www.r-project.org/) using *ggplot2* (Wickham, 2016) package and statistical analysis using *ggpubr*. Machine learning classification was implemented using Sci-kit learn in Python (Pedregosa et al., 2011). The script used for data analysis in R is publicly available as an R-notebook (http://doi.org/10.5281/zenodo.3534239), as well as the Jupyter notebook containing the command lines used for machine learning (http://doi.org/10.5281/zenodo.3534148).

## 3. Results

### 3.1 Extended exposure to heat stress results in a proportional decrease of the rosette size and photosynthetic efficiency

To assess whether high-throughput phenotyping can capture significant alterations in plant physiology caused by exposure to heat stress, we exposed three weeks old Arabidopsis plants to 3 h, 6 h or 9 hours of acute heat stress (45°C), and evaluated the plant size, morphology, temperature, and chlorophyll fluorescence for the subsequent 7 days after stress (DAS) imposition (**Figure 1**). All three heat stress treatments resulted in a significant reduction of the rosette size (**Figure 2B**), and these differences were substantial already 1 h after heat stress treatment. In general, the extended exposure to high temperature increased the effect observed on the rosette size (**Figure 2B**). The applied treatments were sub-lethal to the plants, as only four plants died after 9 h of exposure to heat stress, while other 36 plants that underwent this treatment were able to recover from the stress and produced new green leaves. These results suggest that soil-grown 3-week-old Arabidopsis plants are resilient to acute heat stress and that 6h of heat stress treatment is best at differentiating between WT and heat-sensitive mutant *hsp101*(**Figure 2C**).

**Figure 2.**
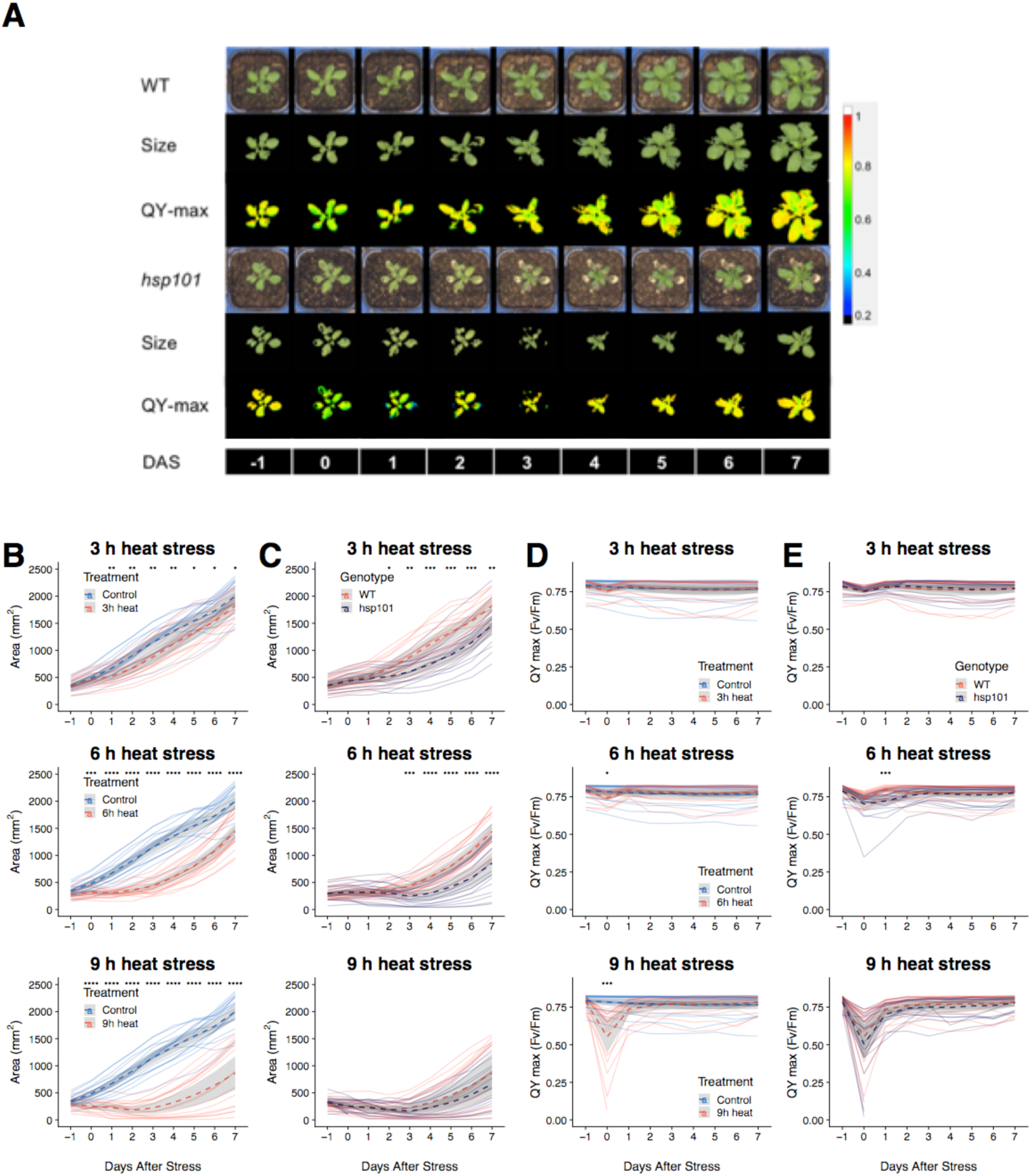
More pronounced reduction of the rosette size with increased length of heat treatment and mutation of HSP101. **(A)** Comparison of rosette size of Col-0 plants grown under control conditions and 3 h, 6 h and 9 h of heat stress (45°C) treatment. **(B)** Comparison of heat-induced changes in the rosette area in WT and *hsp101* for plants exposed to 45 C for 3 h, 6 h, and 9 h. **(C)** Representative RGB, mask and chlorophyll fluorescence images of WT and *hsp101* rosettes of plants underwent 6 h heat stress treatment. The color code depicted at the right represents the maximal quantum yield (QY-max) from blue (low F_v_/F_m_) to red (high F_v_/F_m_). DAS: Days after stress. (**D**) Comparisons of maximal quantum yields of non-stressed WT plants with 3 h, 6 h, and 9 h heat-stressed WT (**E**) Comparisons of maximal quantum yields of 3 h, 6 h, and 9 h heat-stressed WT and *hsp101*. Dashed lines and grey ribbons represent the mean value ± 95 % confidence intervals of different plant groups (n varies between 17 and 20 for individual groups). The significance of the difference in the size of treated and non-treated (A, D), and Col-0 and *hsp101* (B, E) plants each day during the imaging period was determined by Student’s t-test, with p-value below 0.05, 0.01, 0.001 and 0.0001 indicated with *, **, *** and **** respectively.

To further evaluate the changes in relationship to overall heat tolerance, we compared Col-0 (WT) and *hsp101* mutant (**Figure 2A, C**). After heat stress imposition, the *hsp101* plants developed smaller rosettes sizes than WT (**Figure 2C**), while no significant difference in rosette size was observed between WT and hsp101 without heat stress imposition (**Figure S1A**). The significant differences in rosette size were observed between Col-0 and *hsp101* two days after the 3 h heat treatment and three days after the 6 h treatment. No difference between Col-0 and *hsp101* was observed after the 9 h treatment, where the rosette size was reduced in a similar degree (**Figure 2C**).

While plant size broadly reflects overall plant performance, high temperature can also influence plant morphology, such as the petiole elongation and increased leaf angle (Koini et al., 2009; Crawford et al., 2012). We examined the effect of heat stress on rosette morphology and observed that heat stress treatment had a pronounced effect on rosette perimeter, compactness, and slenderness of leaves already one hour after heat stress application (**Figure S2 A**). In addition, these three parameters showed significant differences between heat stress treated Col-0 and *hsp101* lines (**Figure S2 B**). Another morphological change that we observed one day after the heat application was the increase in leaf angle, which occurred only in Col-0 plants but not in *hsp101* (**Figure S2 D**). The change in leaf angle was reflected by the transient increase of rotational mass symmetry in Col-0 at DAS 1, but returned back to levels observed in non-stressed plants 2 days after stress application (**Figure S2 B**).

The decrease in plant photosynthetic efficiency is generally believed to precede developmental changes. Based on the chlorophyll fluorescence we indeed observed a decline in maximum quantum yield (QY max), derived from the measurements of dark-adapted minimum (F_o_, **Figure S3**) and maximum fluorescence (F_m_, **Figure S4**) as (F_m_ – F_o_) /F_m_. QY max indicates the efficiency of PSII photochemistry in the dark-adapted state and reveals the efficiency of electron transport inside PSII. The immediate decline in QY max occurred on the stress day (DAS 0) for both WT and *hsp101* (**Figure 2 D-E**). The WT was able to recover the QY max 1 DAS to levels observed in non-stressed plants, whereas QY max of *hsp101* plants recovered only at 2 DAS (**Figure 2 E**). We also noted the severe reduction of the QY max in *hsp101* at the edges of the rosette (**Figure 2 A**), which preceded the tissue senescence. The earlier reduction in QY max in these areas suggests that change in chlorophyll fluorescence can be used as early indicators of premature senescence.

Exposure to high-temperature was earlier reported to result in an instant increase in minimal (F_0_) and the decrease in maximum fluorescence (F_m_) in various plant species (Schreiber and Armond, 1978; Yamane et al., 1997). The light-curve protocol used to examine the chlorophyll fluorescence, allowed us to study heat-induced changes in light-adapted state at various light intensities (**Figure 1 D**). Heat stress exposure resulted in lower F_m_ measured at 0 DAS (**Figure S4**), while the significant decrease for F_o_ was observed at 1 DAS in WT plants for all three heat stress regimes (**Figure S3 A**). By examining all the directly measured traits at 0 DAS (**Figure 3 A**), we noted a heat-stress induced decrease in light-adapted F_m_ at all studied light intensities. This reduction in maximum fluorescence was more pronounced in *hsp101* plants for the lowest and highest light intensities studied (**Figure 3 B**). The heat stress also resulted in reduced minimal and steady-state fluorescence (F_t_) across studied light intensities at 1 DAS (**Figure 3 C**). While 1 DAS the F_m_ of heat-treated plants was still lower compared to non-treated plants, the difference between WT and *hsp101* at 1 DAS was no longer significant (**Figure 3 D**). These results exemplify the dynamics and rapid recovery of the chlorophyll fluorescence.

**Figure 3.**
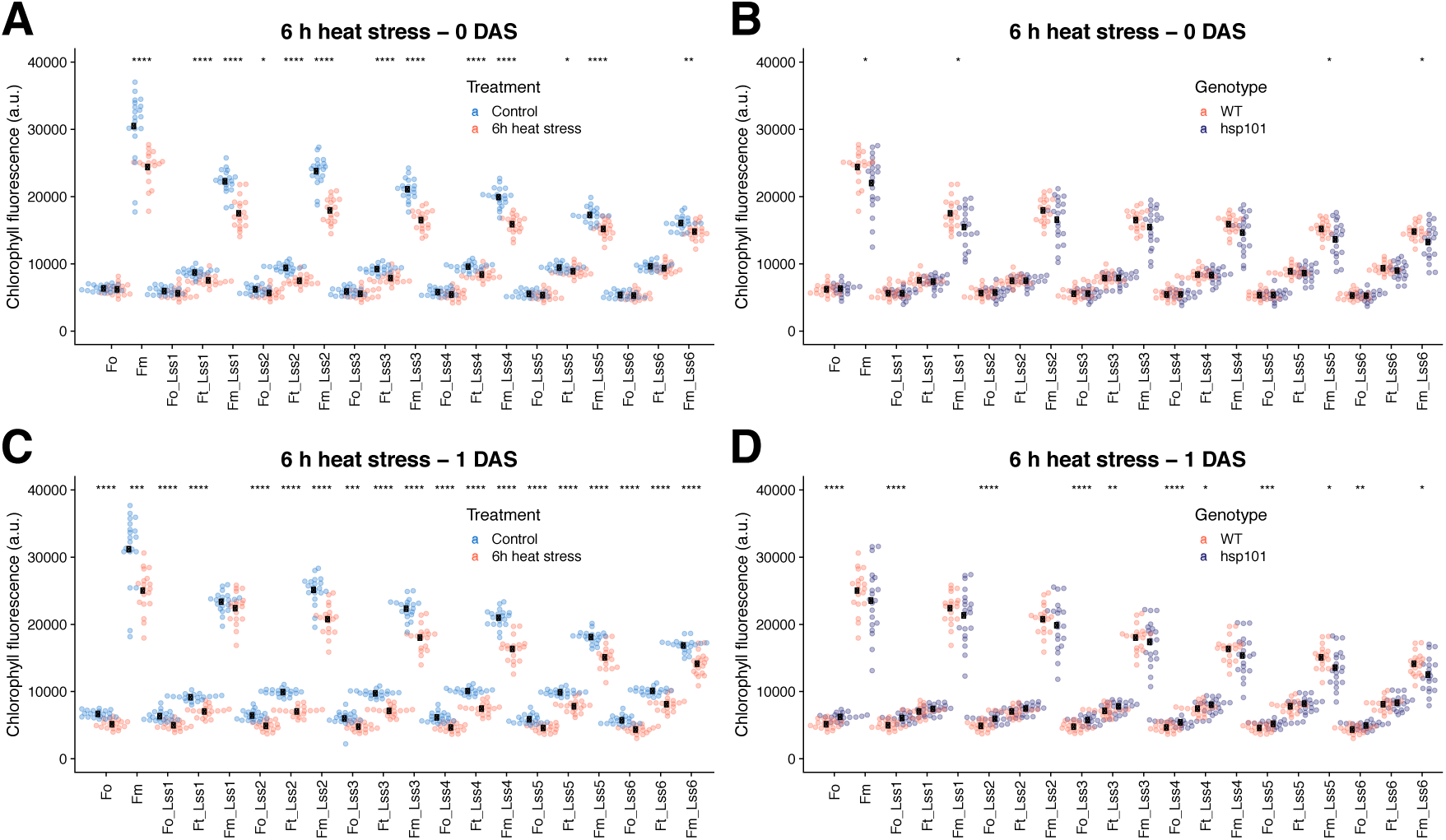
Heat stress-induced changes in chlorophyll fluorescence shows a dynamic profile. The comparison of the directly measured chlorophyll fluorescence traits 1 h after 6 h of heat stress treatment (45°C) application in **(A)** WT compared to WT non-stressed plants and **(B)** WT compared to hsp101 plants exposed to heat stress. The directly measured chlorophyll fluorescence traits were also measured at 1 day after stress (DAS) in **(C)** WT compared to WT non-stressed plants and **(D)** WT compared to hsp101 plants exposed to heat stress. The average of the individual groups are represented by the black dot, while the measurements derived from individual plants are represented using transparent points. The F_o_ and F_m_ indicate minimal and maximal chlorophyll fluorescence measured at dark-adapted state respectively, while F_o_, F_t_ and F_m_ at Lss1 to Lss6 represent minimal, steady-state and maximal chlorophyll fluorescence respectively in light-adapted state at light intensities of 95, 210, 320, 440 and 670 µ mol m^−2^ s^−1^. The significance of the difference in chlorophyll fluorescence of treated and non-treated (A, C), and Col-0 and *hsp101* (B, D) plants for individual parameters was determined by Student’s t-test, with the p-value below 0.05, 0.01, 0.001 and 0.0001 indicated with *, **, *** and **** respectively.

As plant’s cooling capacity is expected to be affected by exposure to high-temperature, **w**e examined differences in leaf temperature recorded from an infrared camera between heat-stressed and non-stressed plants (**Figure S5**). Heat exposure increased leaf temperature across three different heat stress treatments (**Figure S5 A**), no significant differences in leaf temperature were observed between WT and *hsp101* plants (**Figure S5 B**).

### 3.2 Classification model and trait selection to differentiate heat-sensitive and tolerant lines

As the high-throughput phenotyping dataset allows us to examine more than 80 phenotypes, it makes it difficult to rank individual phenotypes to select the best traits which allow clear differentiation of heat-sensitive genotypes. Therefore, we implemented the machine learning to simultaneously explore all the quantitative phenotypes collected within our experiment to differentiate between WT and *hsp101* mutant. We applied logistic regression with lasso regularization to select the most useful traits for classification. To identify the heat stress regime allowing us to differentiate between WT and *hsp101*, we compared the classification performance of phenotypic data from different treatment groups including rosette size, leaf temperature and a subset of top morphological traits (perimeter, the slenderness of leaves, compactness, isotropy, and rotational mass symmetry) and chlorophyll fluorescence parameters measured in dark-adapted state (F_o_, F_m_, and QY max). The accuracies of model prediction (**Table 1**) were modest, with the highest accuracy of 68.2% for the 3 h heat treatment. The predictions calculated for 9 h heat stress treatment was lower than for the non-stressed plants, implying that 9 h heat stress treatment was too severe and not suited to differentiate phenotypic differences between the genotypes. When we included all phenotypic traits in the model, including the chlorophyll fluorescence measured in light adapted state, we observed improved classification accuracies for all groups (**Table 1)**. In line with our earlier analysis (**Figure 2**), the phenotypes recorded for plants treated with 6 h of heat stress provided the highest accuracy (81.5%) in differentiating between two genotypes. The top five contributing parameters to the classification model were related to temperature and morphology (compactness, isotropy, slenderness of leaves and perimeter), suggesting that variability in morphological changes were the most useful genotypic indicators during the overall imaging period. The chlorophyll fluorescence parameters determined to be the most indicative of the genotype are direct measurements of the chlorophyll fluorescence (F_t_’, F_o_, F_o_’) and the photochemical quenching of the chlorophyll fluorescence at the light-adapted state (F_q_’). While some parameters were classified as good genotype indicators under all treatments, e.g. leaf temperature, we noticed that photochemical quenching at the second to highest light intensity (F_q_ Lss5) was unique to the 3 and 6 h of heat stress treatment.

**Table 1.** Logistic regression classification accuracies to determine Col-0 and *hsp101* plants under different treatments. SOL – Slenderness of leaves, RMS – Rotational mass symmetry. F_o_ – minimal fluorescence in dark-adapted state, F_o_’ – minimal fluorescence in light-adapted state, F_t_’ – steady-state fluorescence in the light-adapted state, and F_q_ – photochemical quenching. Lss indicates the light intensity at which individual chlorophyll fluorescence traits were determined (see the material & methods section).

| Treatment | Control | 3 h HS | 6 h HS | 9 h HS |
| --- | --- | --- | --- | --- |
| Accuracy (Subset of traits) | 61.1% | 68.2% | 65.2% | 58.4% |
| Accuracy (all traits) | 67.9% | 79.0% | 81.5% | 65.7% |
| Top 10 traits | Temperature<br>SOL<br>$F_t'$ Lss2<br>$F_t'$ Lss4<br>$F_t'$ Lss3<br>$F_t'$ Lss6<br>$F_q'$ Lss3<br>$F_q'$ Lss6<br>$F_o'$ Lss3<br>$F_q'$ Lss2 | RMS<br>Isotropy<br>SOL<br>$F_o'$ Lss2<br>$F_o$<br>$F_t'$ Lss5<br>$F_q'$ Lss5<br>$F_o'$ Lss3<br>$F_t'$ Lss4<br>$F_t'$ Lss3 | Compactness<br>Isotropy<br>Temperature<br>SOL<br>Perimeter<br>$F_q'$ Lss5<br>$F_t'$ Lss4<br>$F_t'$ Lss3<br>$F_v'$ Lss4<br>$F_t'$ Lss2 | RMS<br>Temperature<br>SOL<br>$F_t'$ Lss5<br>$F_o'$ Lss5<br>$F_o$<br>$F_t'$ Lss1<br>$F_t'$ Lss6<br>$F_t'$ Lss2<br>$F_q'$ Lss6 |

### 3.3 Decline in photochemical quenching as an early indicator of heat stress susceptibility

As photochemical quenching (F_q_) was a unique trait that was associated with differentiation between the WT and heat-sensitive *hsp101* mutant, we examined the heat-induced changes in F_q_ throughout the duration of the experiment (**Figure 4**). The heat exposure resulted in an immediate reduction of F_q_ across all heat stress regimes, and recovery to the F_q_ levels observed for the non-treated plants within 1, 2 or 3 DAS for WT plants exposed to heat stress for 3, 6 and 9 h respectively (**Figure 4 A**). The heat-sensitive *hsp101* mutant showed an even more severe decrease in F_q_ immediately after heat stress exposure in plants exposed to heat stress for 3 and 6 hours (**Figure 4 B**). While we also observed a reduction in the non-photochemical quenching measured at the same light intensity (Lss5) for the WT plants exposed to heat stress (**Figure 4 C)**, the differences between WT and *hsp101* were less pronounced (**Figure 4 D**). The F_q_ measured at other light intensities showed similar trends (**Figure S6**), although the effect of the heat stress on F_q_ differed depending on the light intensity at which the chlorophyll fluorescence was studied.

**Figure 4.**
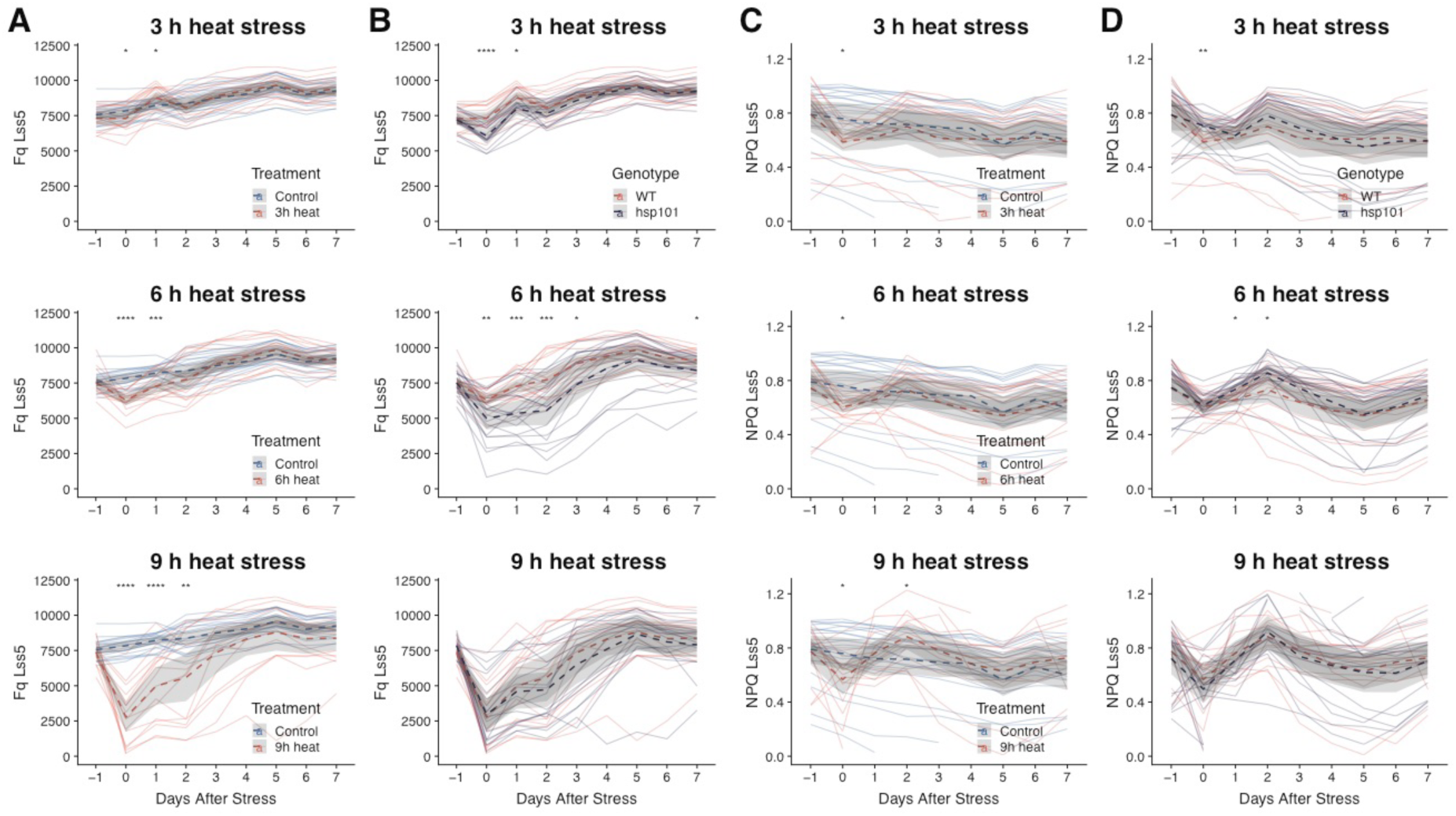
Heat stress transiently reduces photochemical and non-photochemical quenching. **(A)** Comparison of photochemical quenching (F_q_) of Col-0 plants grown under control conditions and 3 h, 6 h and 9 h of heat stress (45°C) treatment. **(B)** Comparison of heat-induced changes in F_q_ in WT and *hsp101* for plants exposed to 45°C for 3 h, 6 h, and 9 h. (**D**) Comparisons of non-photochemical quenching (NPQ) of non-stressed WT plants with 3 h, 6 h, and 9 h heat-stressed WT (**E**) Comparisons of NPQ of 3 h, 6 h and 9 h heat-stressed WT and *hsp101*. Dashed lines and grey ribbons represent the mean value ± 95 % confidence intervals of different plant groups (n varies between 17 and 20 for individual groups). The significance of the difference in the size of treated and non-treated (A, C), and Col-0 and *hsp101* (B, D) plants each day during the imaging period was determined by Student’s t-test, with p-value below 0.05, 0.01, 0.001 and 0.0001 indicated with *, **, *** and **** respectively.

In order to examine whether the heat stress-induced reduction in F_q_ was corresponding to the plant’s performance at the later stage of the experiment, we examined the correlations for individual plants between F_q_ and rosette area measured at individual DAS (**Figure 5**). For plants that did not undergo the heat stress treatment, we observed a correlation between F_q_ and area across all time points (**Figure 5 A)**, however, the correlation within the traits (e.g. area at different DAS) was higher than between the traits. Interestingly, for the plants exposed to heat stress for 3 and 6 h, the F_q_ measured at 0 and 1 DAS shows strong correlations with the rosette area measured throughout the experiment (**Figure 5 B-C**). This suggests that the maintenance of F_q_ could be used as an indicator of plant performance. We also examined the correlations throughout the time between other traits that ranked high in logistic regression classification (**Table 1**) and the rosette area. The heat stress-induced changes in isotropy were negatively correlated with rosette area at earlier time points (DAS 2-4, **Figure S7**). The rosette compactness at 1 DAS also exhibited a small negative correlation with the rosette area at later stages (5-7 DAS) (**Figure S8**). The slenderness of leaves (SOL) scored at early timepoints after stress was not correlated with the plant area at 3 or 6 h of heat stress, and at 9 h the rosette area scored at earlier time points was negatively correlated with SOL, suggesting that observed changes in SOL are rather a consequence of reduced rosette area, rather than the cause (**Figure S9**).

**Figure 5.**
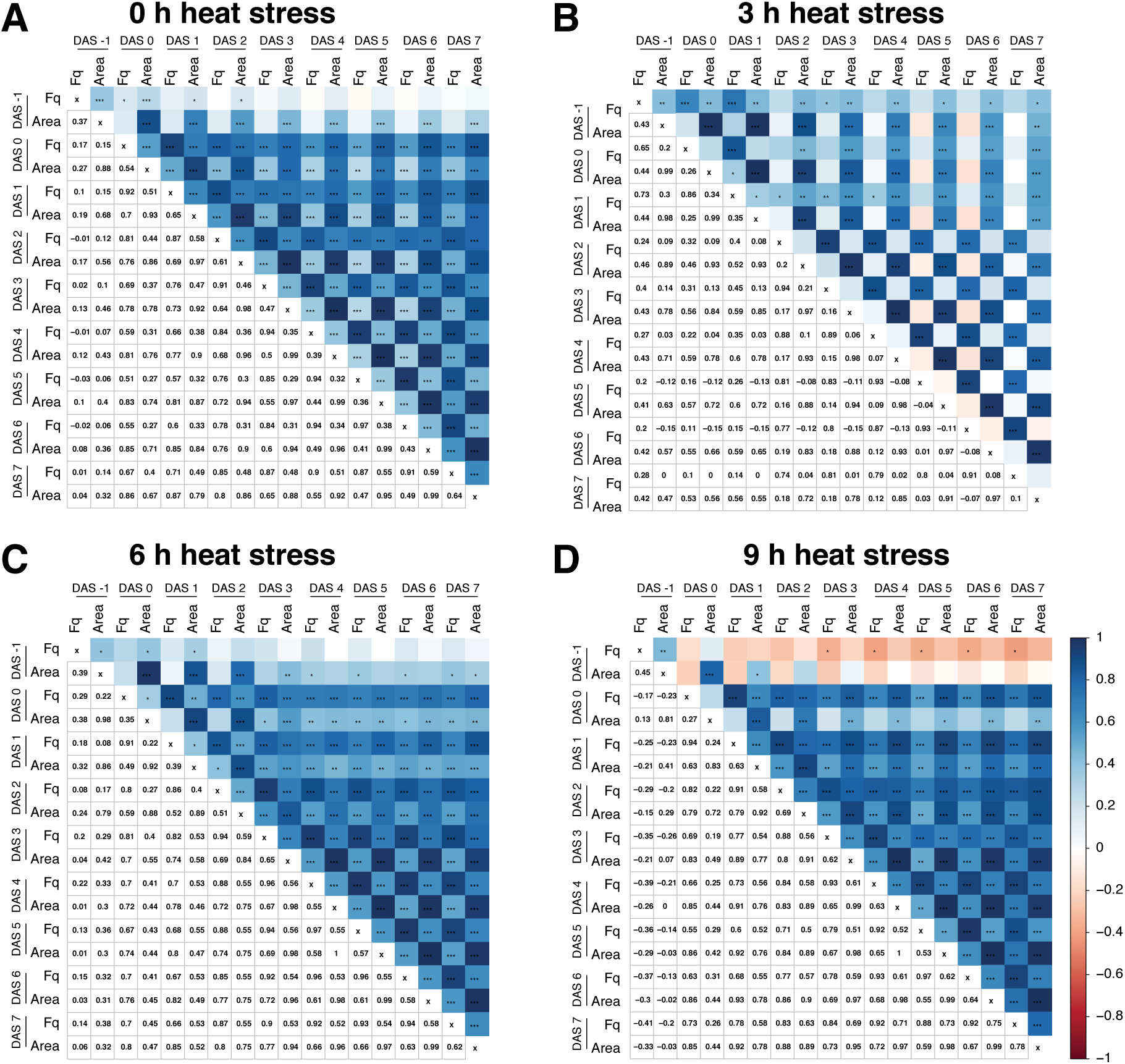
Early changes in photochemical quenching correspond to a larger rosette area. The correlation matrix between photochemical quenching (F_q_) and rosette area scored at various days after stress (DAS) application for the plants **(A)** not-exposed to heat stress **(B)** exposed to 3 h, **(C)** 6 h or **(D)** 9 h of heat stress (45 °C). The positive correlation coefficients are indicated with blue, while negative correlation coefficients with red in the upper part of the correlogram. The Pearson correlation coefficient values are listed as numbers in the lower part of each correlogram. The p-value for each correlation pair, as calculated per t-test, are indicated in the upper part of each correlogram with *, **, *** and **** indicating the p-value below 0.05, 0.01, 0.001 and 0.0001 respectively.

As correlations between rosette area and compactness or isotropy were only weakly significant (0.01 < p-value < 0.05), and the trends were only observed in plants exposed to heat stress for 3 h, the correlation between early changes in F_q_ and rosette area size suggests that maintenance of F_q_ might be causal to plant performance at a later stage. To examine these correlations in more details, the F_q_ at 0 and 1 DAS was plotted for individual heat stress regime and the correlation between the rosette area in individual genotypes was examined (**Figure 6**). The correlation between F_q_ and area, when both traits are scored at 0 DAS, is weak and non-significant in most cases (**Figure 6 A**). However, when we examined the correlation between F_q_ scored at 0 DAS with rosette area at later timepoints, the correlation coefficients increased for the heat-stress treated samples (**Figure 6 B, Figure S10**). Interestingly, across the increasing heat stress exposure, the correlations between F_q_ at DAS 0 and rosette area at later DAS increased more decidedly for the heat susceptible *hsp101*, suggesting that maintenance of photochemical quenching had even higher importance for this heat susceptible genotype. Similar correlations were for F_q_ measured at 1 DAS, where the correlations between F_q_ and rosette area measured both at 1 DAS were relatively weak (**Figure 6 C**), and increase when the rosette area is measured at later time points (**Figure 6 D, Figure S11**). The F_q_ measured at 7 DAS also shows significant correlations with the rosette area measured at the same time (**Figure 6 E**), however the correlation coefficients are lower and the p-values are higher than for the correlations between early changes in F_q_ and later size of the rosette area. These results are strongly suggestive that photochemical quenching might be an important component of heat stress tolerance, and that maintenance of F_q_ is particularly important in the heat susceptible lines.

**Figure 6.**
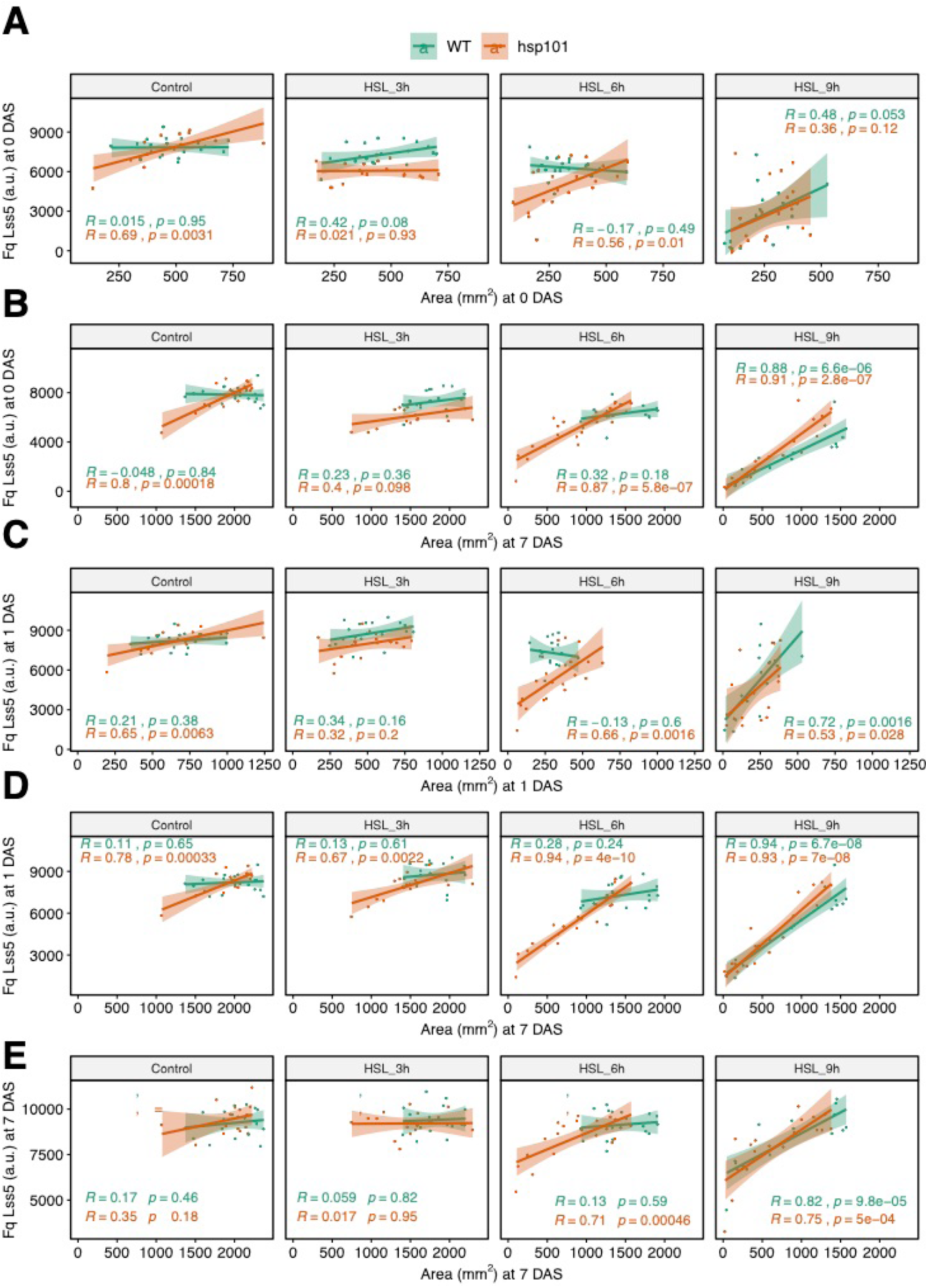
Heat-stress induced reduction in photochemical quenching indicates heat susceptibility. The correlation between photochemical quenching (F_q_) and rosette area was examined for plants not-exposed to heat stress, and plants exposed to 3 h, 6 h or 9 h of heat stress (45 °C). The correlation was examined between Fq scored at 0 days after stress (DAS) imposition and **(A)** rosette area at 0 DAS and **(B)** 7 DAS, between Fq scored at 1 DAS and **(C)** rosette area at 1 DAS and **(D)** 7 DAS, as well as **(E)** F_q_ scored at 7 DAS and rosette area at 7 DAS. The correlation coefficients (R) for individual genotypes (Col-0 WT and *hsp101*) as well as the corresponding p-values as calculated per Pearson correlation t-test are indicated with different colors. The lines in each graph represent the regression line for each treatment and genotype combination. The individual plants are represented by individual dots.

## 4. Discussion

Increasing temperature is one of the most important environmental factors affecting the agricultural productivity worldwide. Improving our understanding of the mechanisms underlying plant heat stress responses will facilitate the development of technologies and breeding strategies for improving plant thermotolerance. In this manuscript, we show an example of how high-throughput phenotyping can be used to screen for heat tolerance related traits, providing more insight into the physiological processes contributing to thermotolerance.

By studying the heat-induced changes in plant size, morphology, temperature and chlorophyll fluorescence we identified a set of phenotypes like leaf temperature, maximum quantum yield, and slenderness of leaves that show immediate response to heat stress. By evaluating three different heat stress regimes (3, 6 and 9 h at 45°C, **Figure 1**), we identified which trends were observed across all treatments. By including a heat-sensitive *hsp101* mutant, we were able to distinguish which phenotypes were informative in distinguishing between WT and heat sensitive lines (**Table 1**). We observed that *hsp101* plants showed a more severe decrease in plant growth and quantum yield compared to the WT plants, and that the difference between genotypes were most evident across the plants exposed to heat stress for 6 h (**Figure 2)**. The rapid change in quantum yield seems to be specific to heat stress, as Arabidopsis plants exposed to salt stress did not show reduced photosynthetic yield with similar length of the experiment (Awlia et al., 2016), while drought decreased quantum yield only at the later stage of stress exposure (Jansen et al., 2009). While the changes in quantum yield and photochemical quenching was observed immediately after the heat stress, the heat stress also affected rosette morphology at later time points, including the reduction in the slenderness of leaves, compactness and increased rosette isotropy (**Figure S2**). Such changes in the rosette morphology were less apparent under salt stress (Awlia et al., 2016). These differences between heat, drought and salt reflect the nature of abiotic stress, with heat stress exposure being the most acute, while the severity of salt and drought stress increases gradually over longer periods of time. The observed differences highlight the importance of selecting the physiologically relevant levels of stress, considering the differences in the nature of individual abiotic stress that the plants are exposed to the environment. Summarizing, results presented in this study showed that plant physiological responses to high temperature are complex and temporal in their nature, with short term changes being captured best with chlorophyll fluorescence (F_m_, F_0_, and QY max, F_q_) and leaf temperature, while heat-induced changes in rosette morphology (rosette area, perimeter, compactness, and slenderness of leaves) are observed at extended time after the application of heat stress. While the majority of the heat stress studies focuses on survival, we think that using the combination of these parameters and screening them at a continuous time-window will provide a better understanding of processes underlying heat tolerance.

In this study, we observed that while heat stress exposure increased the leaf angle in Col-0, this response was absent in *hsp101* mutant lines (**Figure S3**). While the response is obvious to the human eye, it proved difficult to detect it from the available morphology parameters, which use only top view images. The increased leaf angle was earlier suggested to have an adaptive advantage under high temperatures, increasing the cooling capacity of the plant (Crawford et al., 2012; Bridge et al., 2013). As we did not observe any significant differences in leaf temperature between WT and *hsp101* mutants (**Figure S5**), the cooling advantage of this response is yet to be demonstrated. Nevertheless, the *hsp101* showed a greater reduction in maximum quantum yield compared to WT (**Figure 2**), and a lower decrease in photochemical and non-photochemical quenching (**Figure 4**). However, as the change in leaf angle was only observed 1 DAS, it is unlikely that this response to causal to the decreased photosynthetic efficiency. How an increase in leaf angle and photosynthetic quenching mechanisms are orchestrated by HSP101 is beyond the scope of this study. If the leaf angle is to be used for future assays, it is our suggestion to include the 3D measurements of the rosettes using the 3D scanner technology and/or side-view image.

While high throughput phenotyping provides more information on plant performance using the non-destructive measurements, the number of the collected direct and derived measurements (**Table S1**) can be overwhelming for effective trait evaluation. Machine learning algorithms can help navigate through the complex dataset. Previous study used a combination of random forest and support vector machine models to identify most distinguishing root traits and cultivar differentiation across European pea cultivars (Zhao et al., 2016). In our study, we used logistic regression to identify the most informative traits which would enable to differentiate between WT and heat sensitive *hsp101* mutant (**Table 1**). The results supported our earlier analysis that 6 h heat treatment is most informative for differentiate between WT and heat sensitive genotype, and highlighted the significance of morphology traits, direct chlorophyll fluorescence measurements as well as photochemical quenching (**Table 1**). These results lead us to investigate changes in photochemical quenching in more detail (**Figure 4**). We observed that maintenance of photochemical quenching was positively correlated with larger rosettes at later timepoints (**Figure 5**) and that this correlation was predominantly found in the plants exposed to heat stress (**Figre 6**). The contributions of non-photochemical quenching to heat stress tolerance (Havaux et al., 1988), as well as other abiotic stress (Flagella et al., 1995) are widely described in earlier literature. The photochemical quenching is related to the redox state of the first electron acceptor of Photosystem II (Schreiber et al., 1986), but how this process is affected by stress and what is its contribution to overall environmental stress tolerance remains unknown. While we want to stress that the correlation does not prove causation, the observed correlation between photochemical quenching and thermotolerance will be an important cornerstone in future research of heat stress responses.

Using kinetic chlorophyll fluorescence assays, such as light curve protocol (Henley, 1993; Rascher et al., 2000) used in this study, provided new insights in dynamic changes to photosynthetic efficiency and heat stress-induced photochemical quenching. We think that the new phenotypic traits, presented in this manuscript will provide better insight and identify novel players contributing to overall plant performance and heat stress tolerance. As processes contributing to overall thermotolerance are complex (Kotak et al., 2007) and can be acquired in various ways (Burke, 2001), the assays focusing on seedling survival or hypocotyl elongation are of limited value. Using more refined phenotypes, such as rosette morphology parameters or traits derived from direct chlorophyll fluorescence measurements will in future studies contribute to discovery of genes and processes which small and transient, but significant, contribution to thermotolerance.

## General

We thank Dr. Andrew Yip for discussion and technical assistance in machine learning modeling, Eunje Kim with pilot experiment, and Growth Facility and KAUST Core Lab staff, Mr. Thomas Hoover, Mr. John Ramer and Mr. Johnard Balangue for assistance with the phenotyping facility.

## Author contributions

G.G., M.A.T. and M.M.J. designed the experiment, G.G. performed the experiment, G.G. and M.M.J analyzed the data and wrote the manuscript, M.A.T. edited the manuscript.

## Supplementary Materials

**Table S1:**
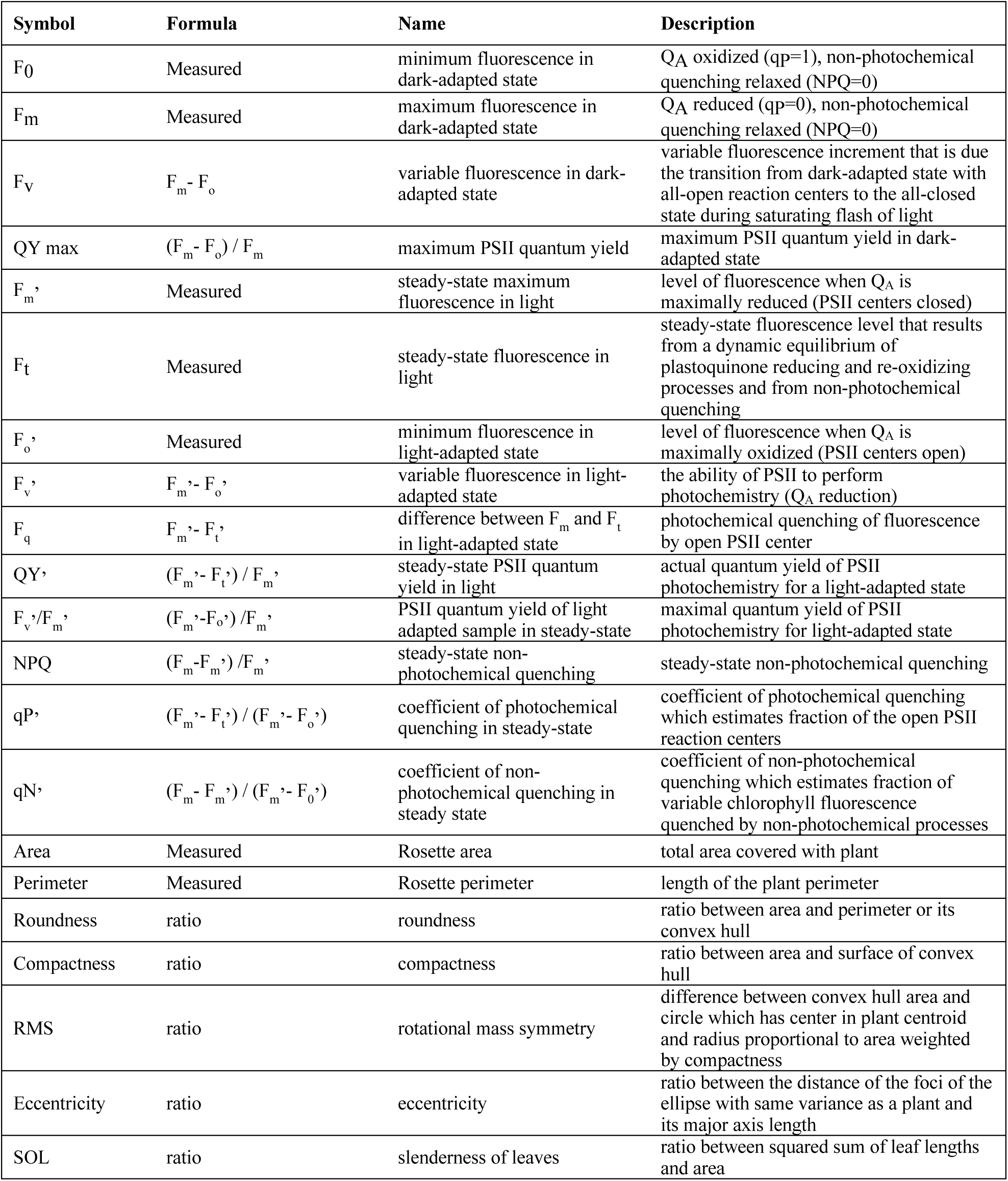
List of direct and indirect chlorophyll fluorescence and morphological parameters used in this study. Q_A_: Quinone A, the primary stable electron acceptor of PSII centers, PSII: Photosystem II, NPQ: non-photochemical quenching

**Figure S1.**
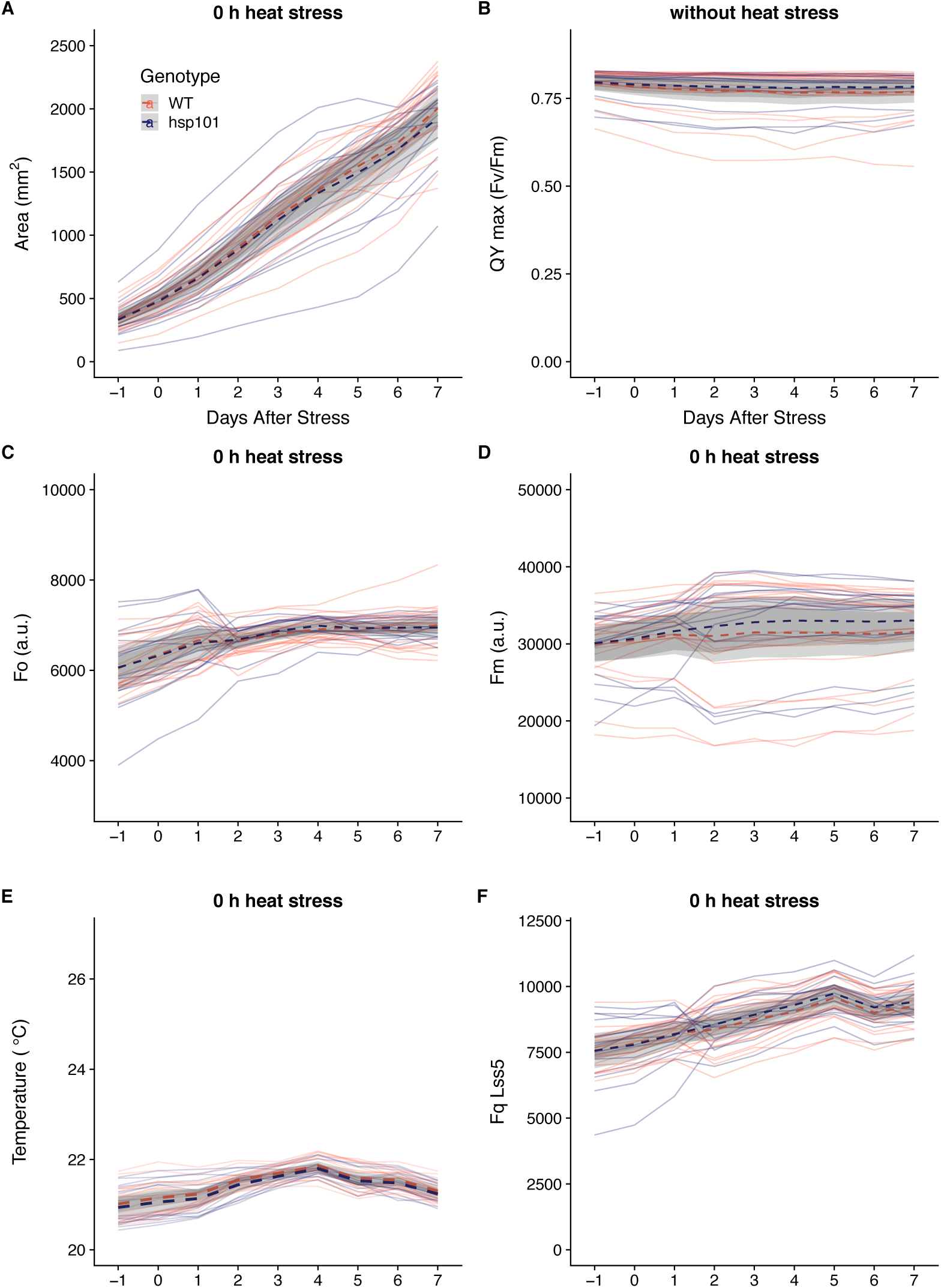
Col-0 and *hsp101* plants are indistinguishable when not exposed to heat stress. Changes in **(A)** rosette area **(B)** maximal quantum yield, **(C)** minimum fluorescence (F_o_), **(D)** maximum fluorescence (F_m_), **(E)** leaf temperature and (F) photochemical quenching (F_q_) were examined in WT and *hsp101* plants grown under control conditions, without heat exposure, during the imaging period of the experiment. Dashed lines and grey ribbons represent the mean values ± 95 % confidence intervals respectively of different plant groups (n = 18 for WT and *hsp101*).

**Figure S2:**
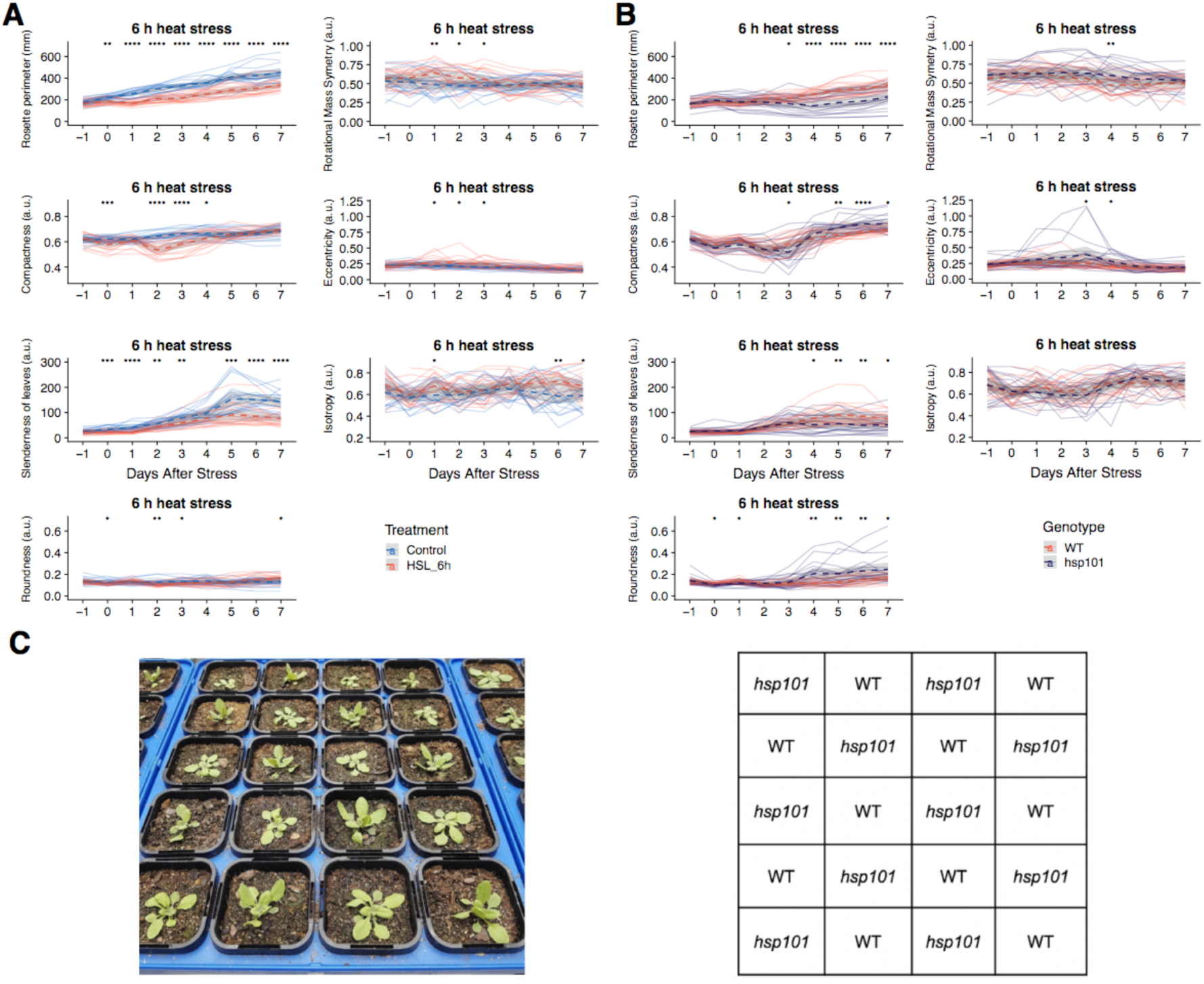
Characterization of heat-induced morphological responses in WT and *hsp101* after 6 h heat treatment. Daily changes in rosette morphology traits were observed in **(A)** non-stressed WT compared to 6 h heat-stressed WT plants, as well as **(B)** and hsp101 plants exposed to heat stress for 6 h. The descriptions of morphology traits are listed in Table S1. Individual lines represent the change in individual plants, while the dash lines and grey ribbons represent the mean values ± 95% confidence intervals of different plant groups (n varies between 19 and 20 plants). Significant difference between heat-stressed and non-stressed WT (A), and Col-0 and hsp101 (B) plants for individual days after stress during the imaging period were determined by Student’s t-test, with p-value below 0.05, 0.01, 0.001 or 0.0001 is indicated with *, **, *** and **** respectively. (**C**) A photo of 6 h heat stress treated WT Col-0 and *hsp101* plants one day after stress application. The position of the genotype is indicated in the tray map on the right panel.

**Figure S3.**
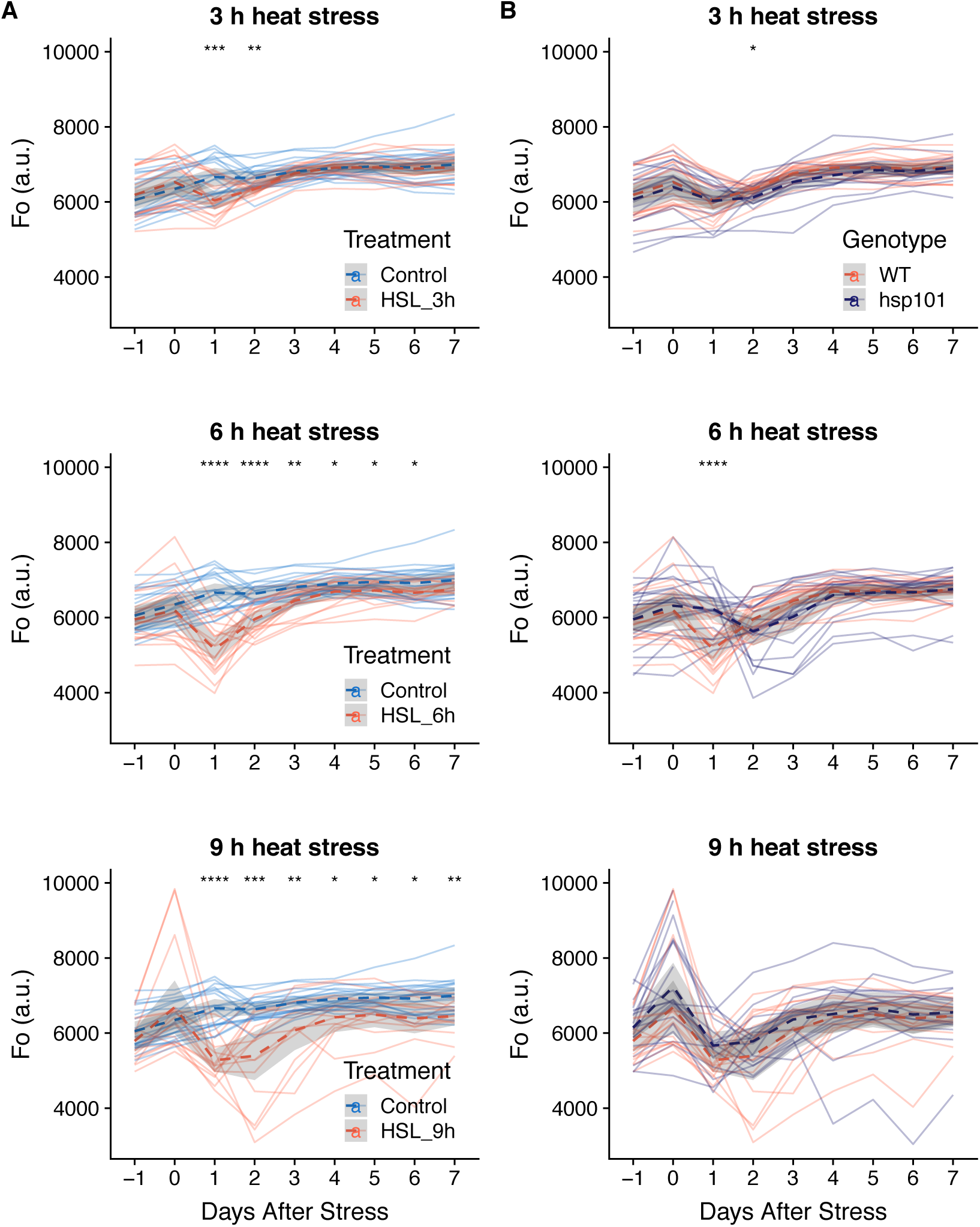
Heat stress induced changes to minimum chlorophyll fluorescence. Daily changes in minimum chlorophyll fluorescence (F_o_) were observed in **(A)** non-stressed WT compared to 6 h heat-stressed WT plants, as well as **(B)** and hsp101 plants exposed to heat stress for 6 h. Individual lines represent the change in individual plants, while the dash lines and grey ribbons represent the mean values ± 95% confidence intervals of different plant groups. Significant difference between heat-stressed and non-stressed WT (A), and Col-0 and *hsp101* (B) plants for individual days after stress during the imaging period were determined by Student’s t-test, with p-value below 0.05, 0.01, 0.001 or 0.0001 is indicated with *, **, *** and **** respectively.

**Figure S4.**
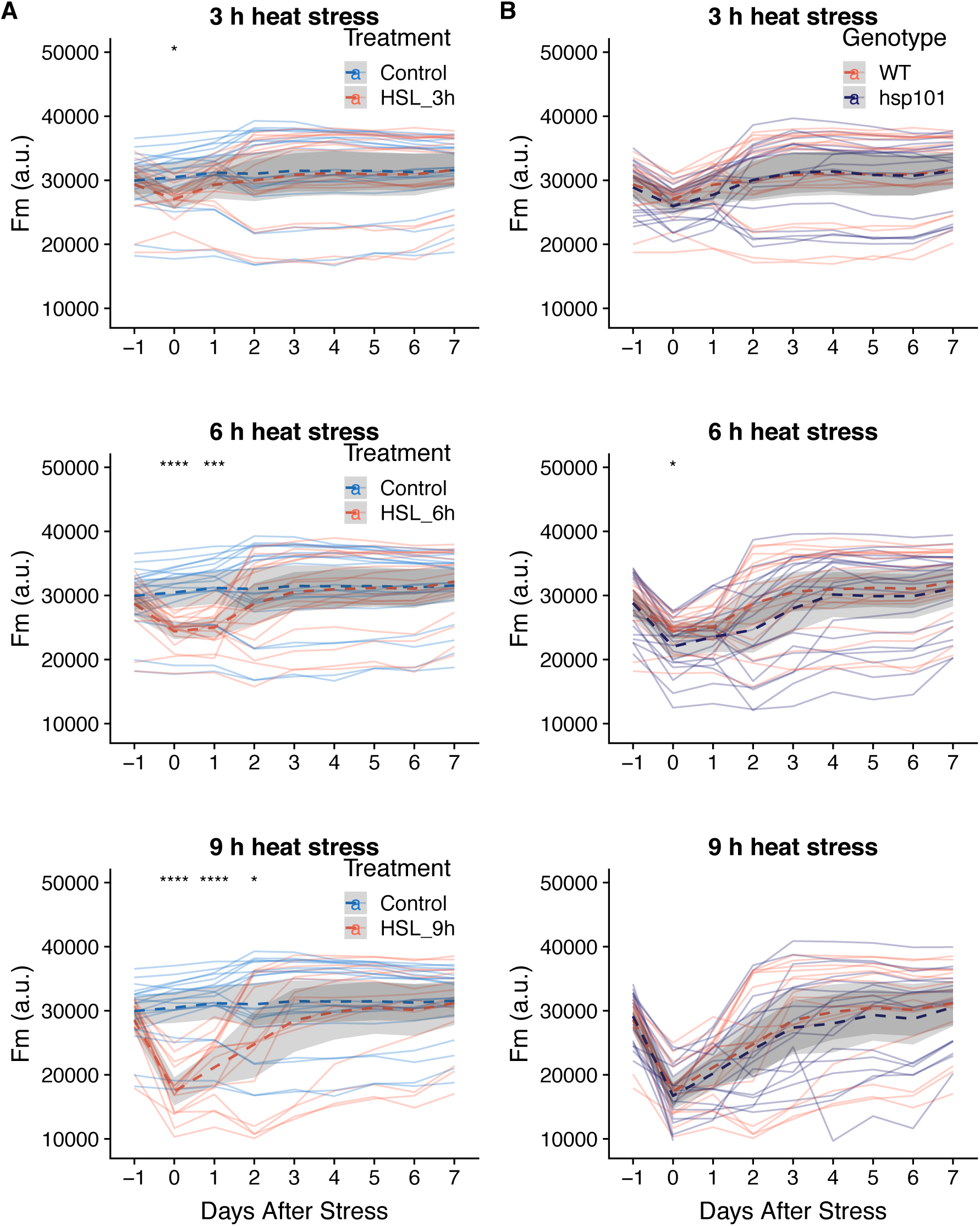
Heat stress induced changes to maximum chlorophyll fluorescence. Daily changes in maximum chlorophyll fluorescence (F_m_) were observed in **(A)** non-stressed WT compared to 6 h heat-stressed WT plants, as well as **(B)** and *hsp101* plants exposed to heat stress for 6 h. Individual lines represent the change in individual plants, while the dash lines and grey ribbons represent the mean values ± 95% confidence intervals of different plant groups. Significant difference between heat-stressed and non-stressed WT (A), and Col-0 and *hsp101* (B) plants for individual days after stress during the imaging period were determined by Student’s t-test, with p-value below 0.05, 0.01, 0.001 or 0.0001 is indicated with *, **, *** and **** respectively.

**Figure S5.**
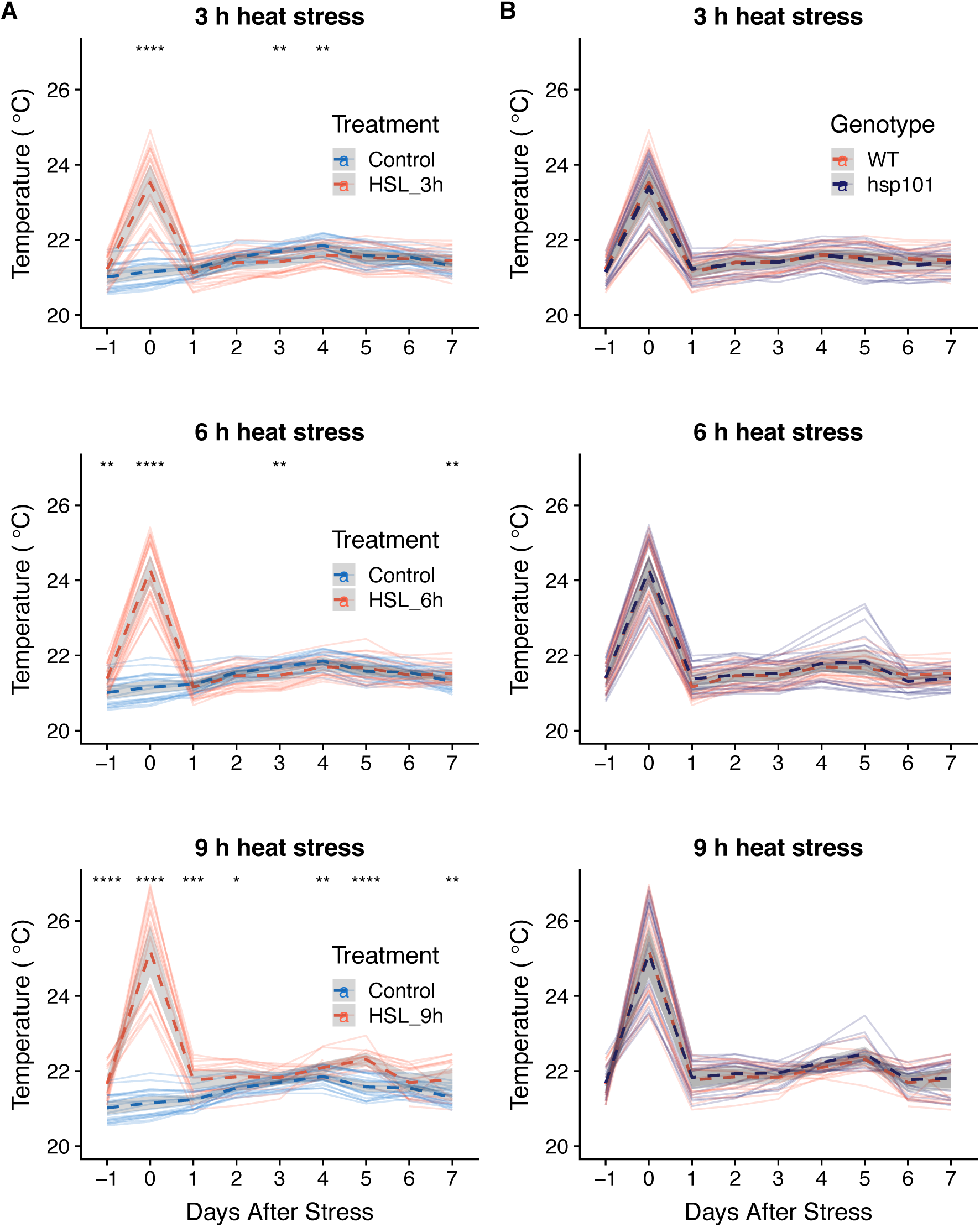
Heat stress induced changes to Leaf temperature. Daily changes in leaf temperature were observed in **(A)** non-stressed WT compared to 6 h heat-stressed WT plants, as well as **(B)** and *hsp101* plants exposed to heat stress for 6 h. Individual lines represent the change in individual plants, while the dash lines and grey ribbons represent the mean values ± 95% confidence intervals of different plant groups. Significant difference between heat-stressed and non-stressed WT (A), and Col-0 and *hsp101* (B) plants for individual days after stress during the imaging period were determined by Student’s t-test, with p-value below 0.05, 0.01, 0.001 or 0.0001 is indicated with *, **, *** and **** respectively.

**Figure S6:**
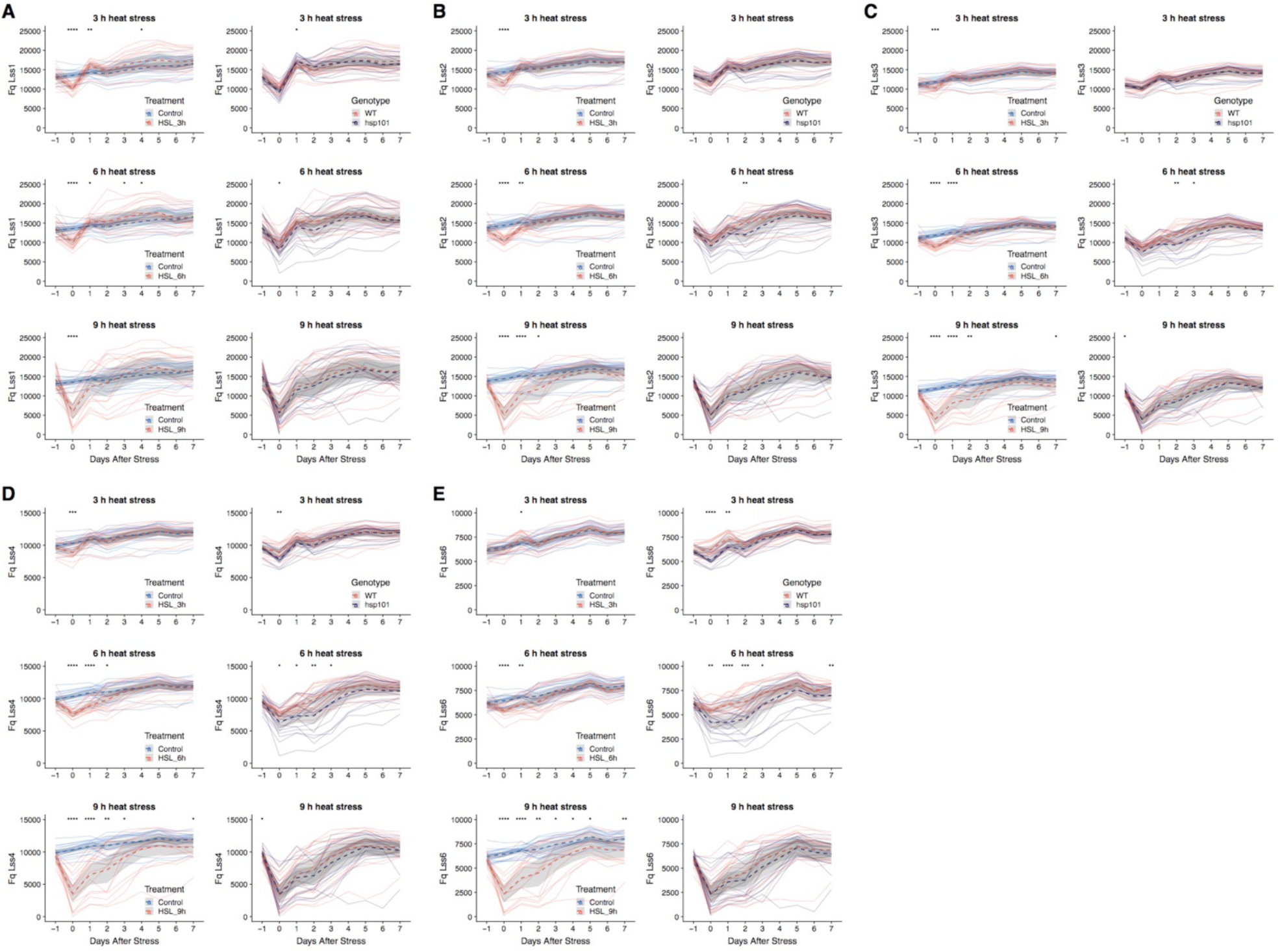
Characterization of heat-induced changes in photochemical quenching in WT and *hsp101*. Daily changes in photochemical quenching (F_q_) observed in non-stressed WT compared to 6 h heat-stressed WT plants, as well as and *hsp101* plants exposed to heat stress for 6 h. The photochemical quenching was calculated based on the light-adapted chlorophyll fluorescence (Table S1) at light intensities of **(A)** 95, (B) 210, (C) 320, (D) 440 and (E) 670 µ mol m^−2^ s^−1^. Individual lines represent the change in individual plants, while the dash lines and grey ribbons represent the mean values ± 95% confidence intervals of different plant groups (n varies between 19 and 20 plants). Significant difference between heat-stressed and non-stressed WT, and Col-0 and *hsp101* plants for individual days after stress during the imaging period were determined by Student’s t-test, with p-value below 0.05, 0.01, 0.001 or 0.0001 is indicated with *, **, *** and **** respectively.

**Figure S7.**
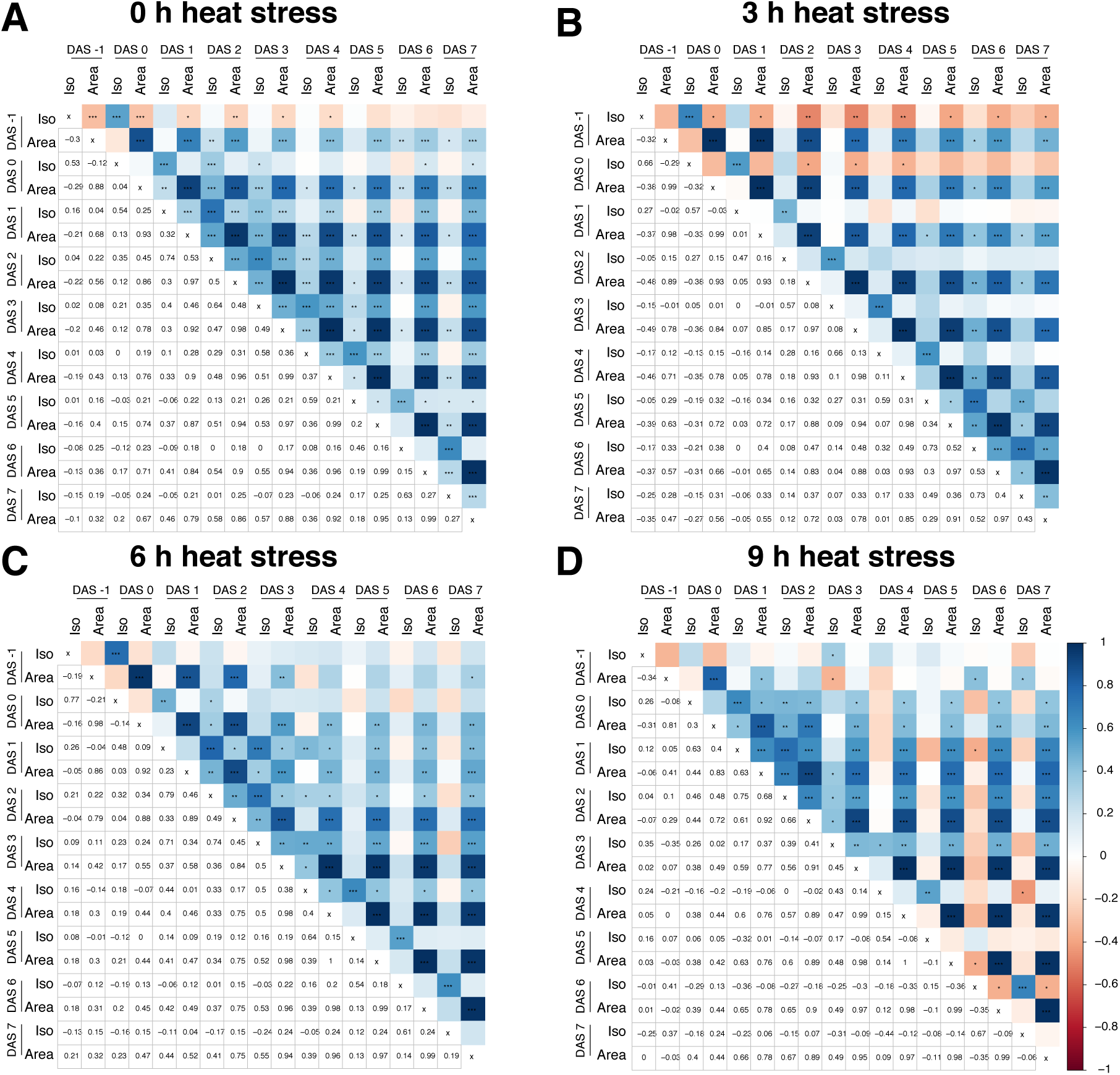
Temporal correlation between heat stress induced changes in rosette isotropy and rosette area. The correlation matrix between rosette isotropy and rosette area scored at various days after stress (DAS) application for the plants **(A)** not-exposed to heat stress **(B)** exposed to 3 h, **(C)** 6 h or **(D)** 9 h of heat stress (45 °C). The positive correlation coefficients are indicated with blue, while negative correlation coefficients with red in the upper part of the correlogram. The correlation coefficient values are listed as numbers in the lower part of each correlogram. The p-value for each correlation pair, as calculated per Pearson correlation t-test, are indicated in the upper part of each correlogram with *, **, *** and **** indicating the p-value below 0.05, 0.01, 0.001 and 0.0001 respectively.

**Figure S8.**
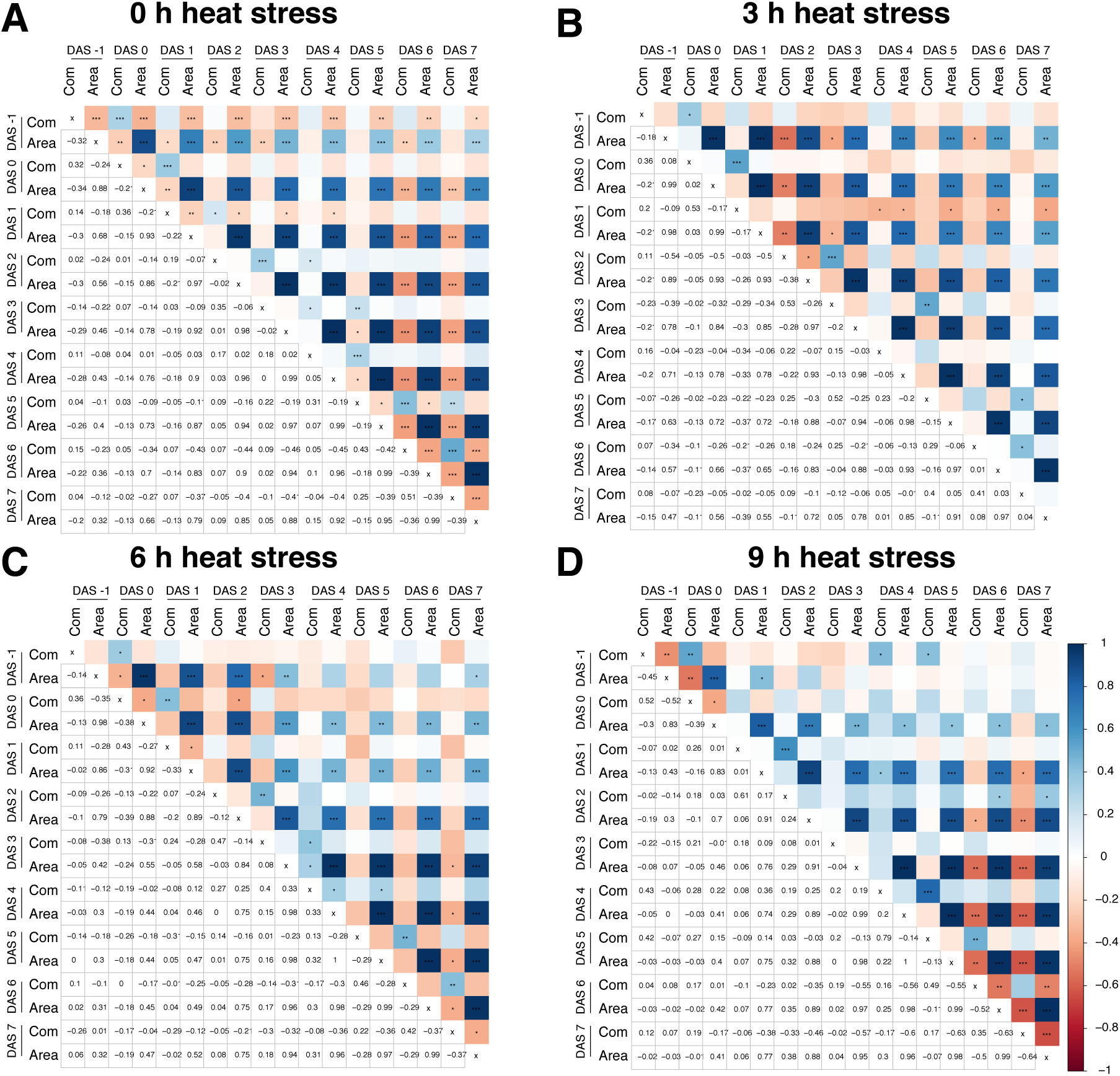
Temporal correlation between heat stress induced changes in rosette compactness and rosette area. The correlation matrix between rosette compactness and rosette area scored at various days after stress (DAS) application for the plants **(A)** not-exposed to heat stress **(B)** exposed to 3 h, **(C)** 6 h or **(D)** 9 h of heat stress (45 °C). The positive correlation coefficients are indicated with blue, while negative correlation coefficients with red in the upper part of the correlogram. The correlation coefficient values are listed as numbers in the lower part of each correlogram. The p-value for each correlation pair, as calculated per Pearson correlation t-test, are indicated in the upper part of each correlogram with *, **, *** and **** indicating the p-value below 0.05, 0.01, 0.001 and 0.0001 respectively.

**Figure S9.**
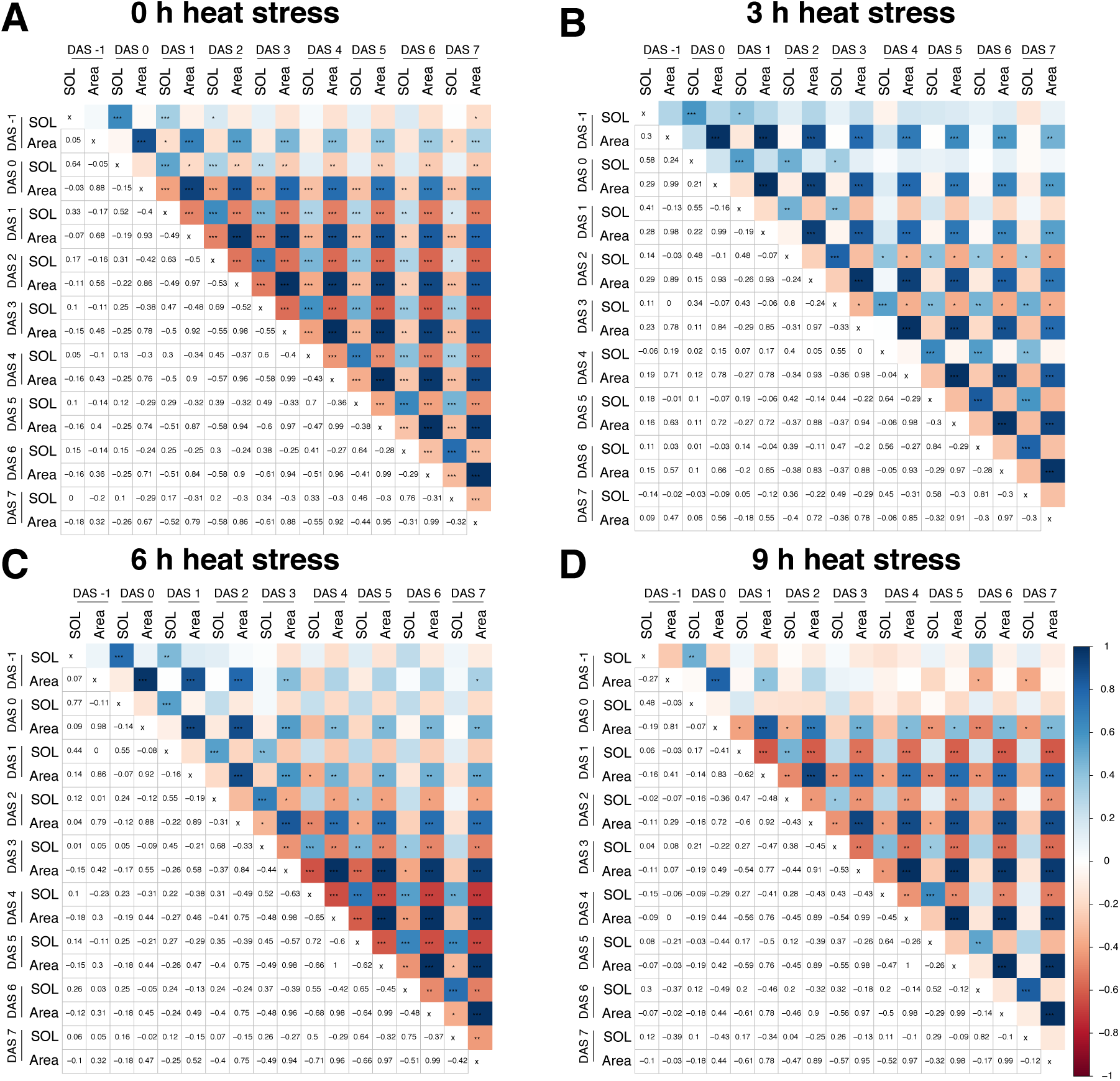
Temporal correlation between heat stress induced changes in slenderness of leaves and rosette area. The correlation matrix between slenderness of leaves and rosette area scored at various days after stress (DAS) application for the plants **(A)** not-exposed to heat stress **(B)** exposed to 3 h, **(C)** 6 h or **(D)** 9 h of heat stress (45 °C). The positive correlation coefficients are indicated with blue, while negative correlation coefficients with red in the upper part of the correlogram. The correlation coefficient values are listed as numbers in the lower part of each correlogram. The p-value for each correlation pair, as calculated per Pearson correlation t-test, are indicated in the upper part of each correlogram with *, **, *** and **** indicating the p-value below 0.05, 0.01, 0.001 and 0.0001 respectively.

**Figure S10.**
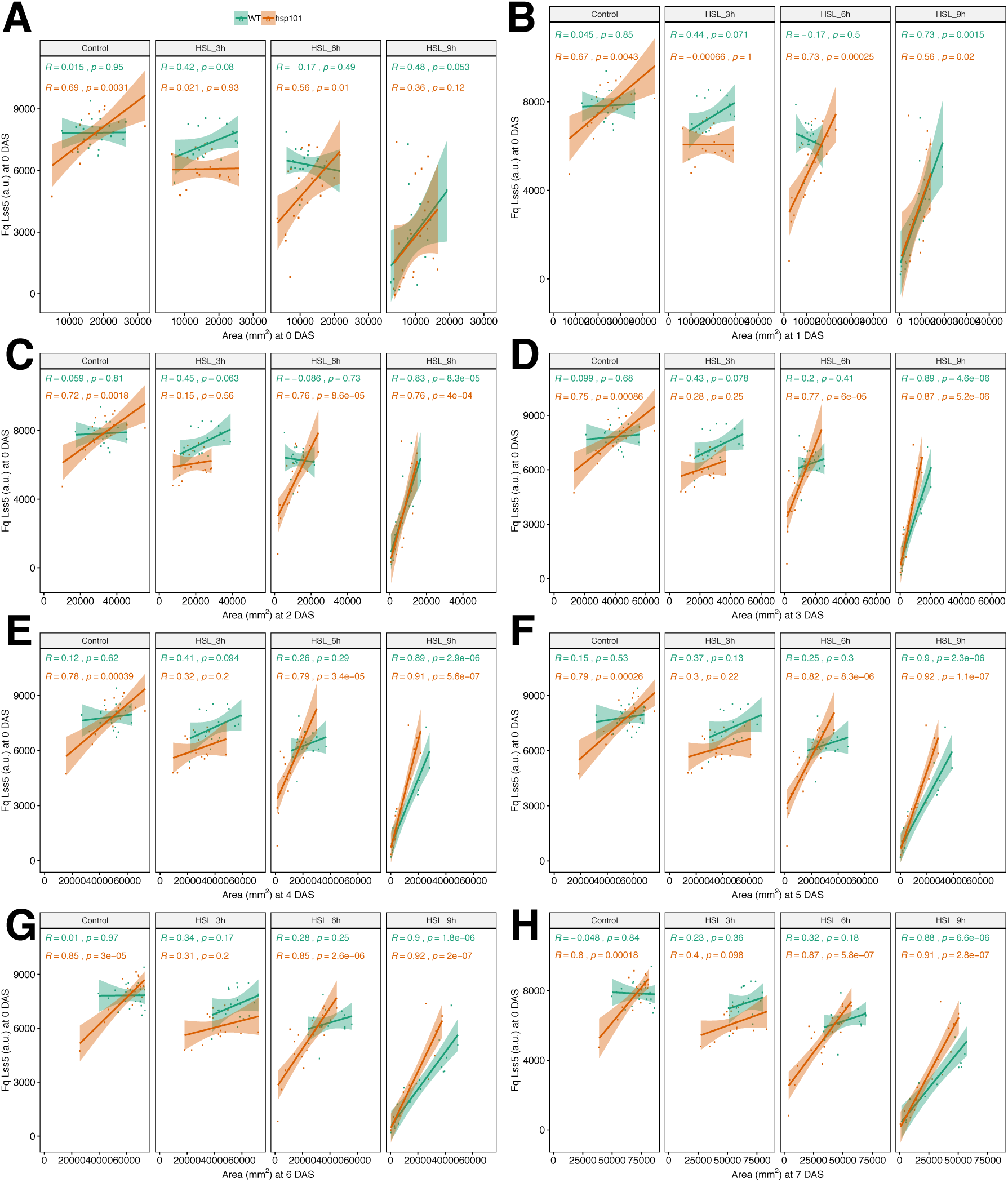
Heat-stress induced reduction in Fq at 0 DAS indicates heat susceptibility. The correlation between photochemical quenching (Fq) and rosette area was examined for plants not-exposed to heat stress, and plants exposed to 3 h, 6 h or 9 h of heat stress (45 °C). The correlation was examined between F_q_ scored at 0 days after stress (DAS) imposition and rosette area at **(A)** 0, **(B)** 1, **(C)** 2, **(D)** 3, **(E)** 4, **(F)** 5, **(G)** 6 and **(H)** 7 DAS. The correlation coefficients (R) for individual genotypes (Col-0 WT and *hsp101*) as well as the corresponding p-values as calculated per Pearson correlation t-test are indicated with different colors. The lines in each graph represent the regression line for each treatment and genotype combination. The individual plants are represented by individual dots.

**Figure S11.**
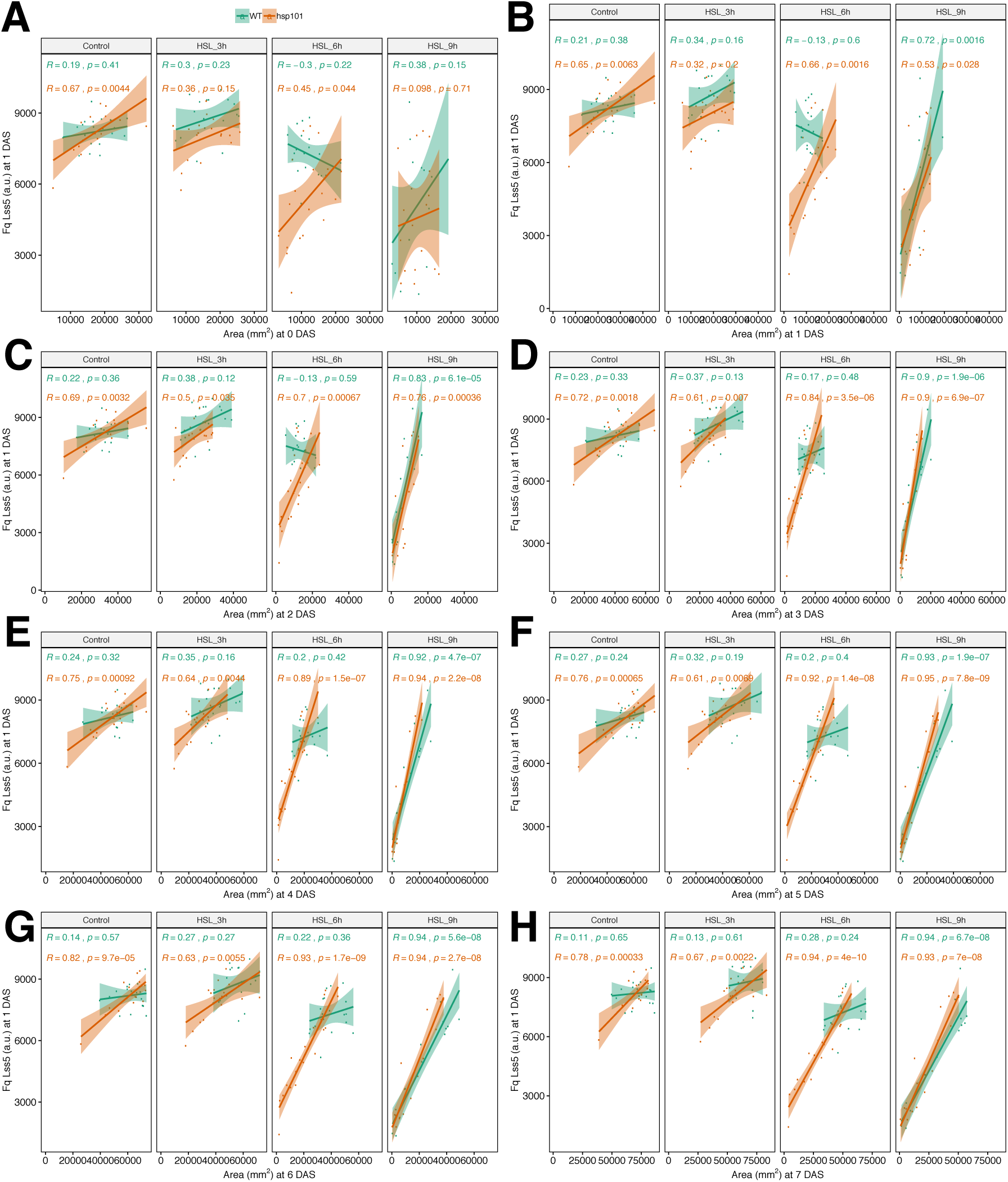
Heat-stress induced reduction in Fq at 1 DAS indicates heat susceptibility. The correlation between photochemical quenching (Fq) and rosette area was examined for plants not-exposed to heat stress, and plants exposed to 3 h, 6 h or 9 h of heat stress (45 C). The correlation was examined between F_q_ scored at 1 days after stress (DAS) imposition and rosette area at **(A)** 0, **(B)** 1, **(C)** 2, **(D)** 3, **(E)** 4, **(F)** 5, **(G)** 6 and **(H)** 7 DAS. The correlation coefficients (R) for individual genotypes (Col-0 WT and *hsp101*) as well as the corresponding p-values as calculated per Pearson correlation t-test are indicated with different colors. The lines in each graph represent the regression line for each treatment and genotype combination. The individual plants are represented by individual dots.

