## Supplementary figures and images for "The use of high throughput phenotyping for assessment of heat stress-induced changes in Arabidopsis"

### Figure S1

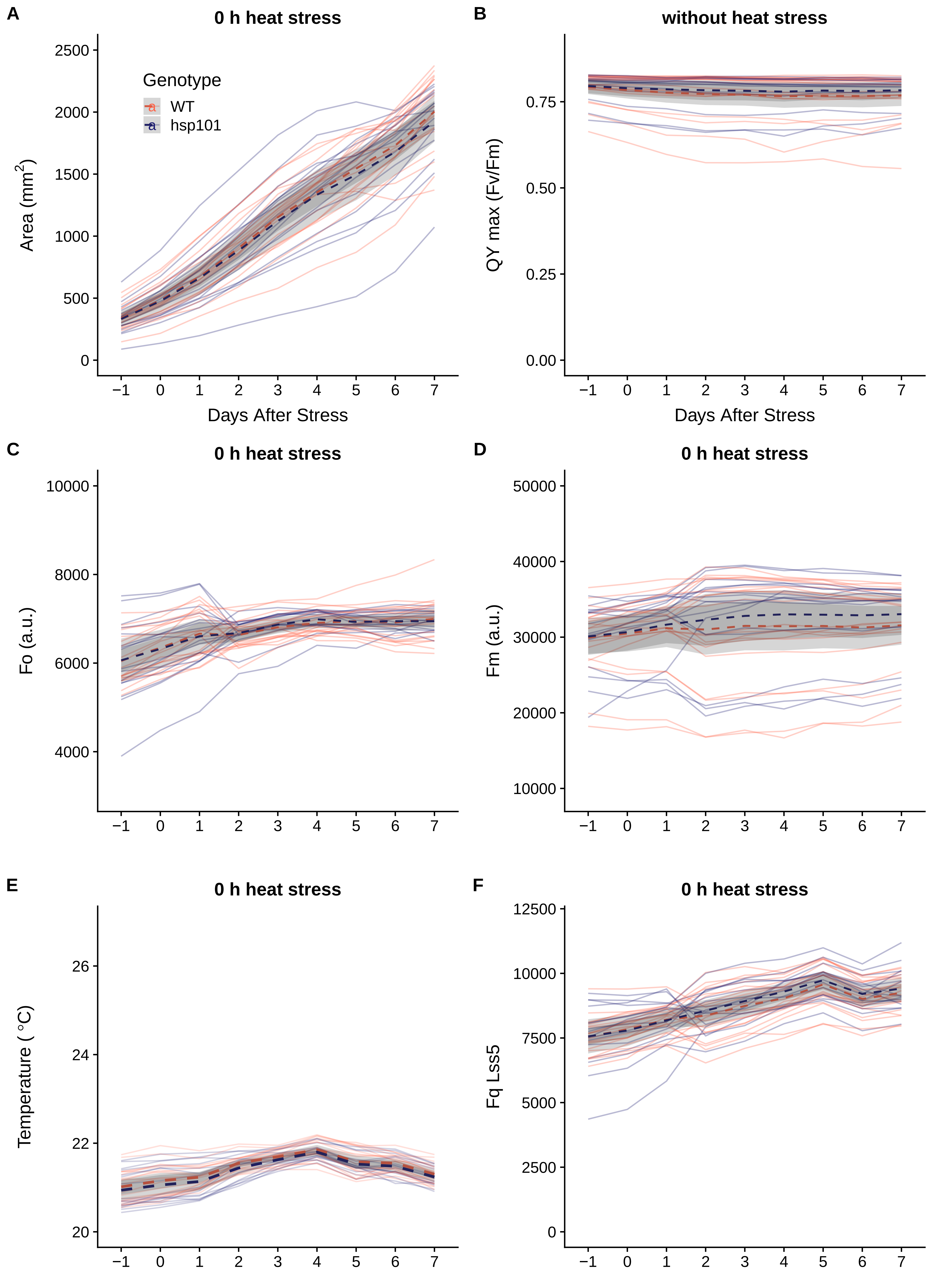

### Figure S2

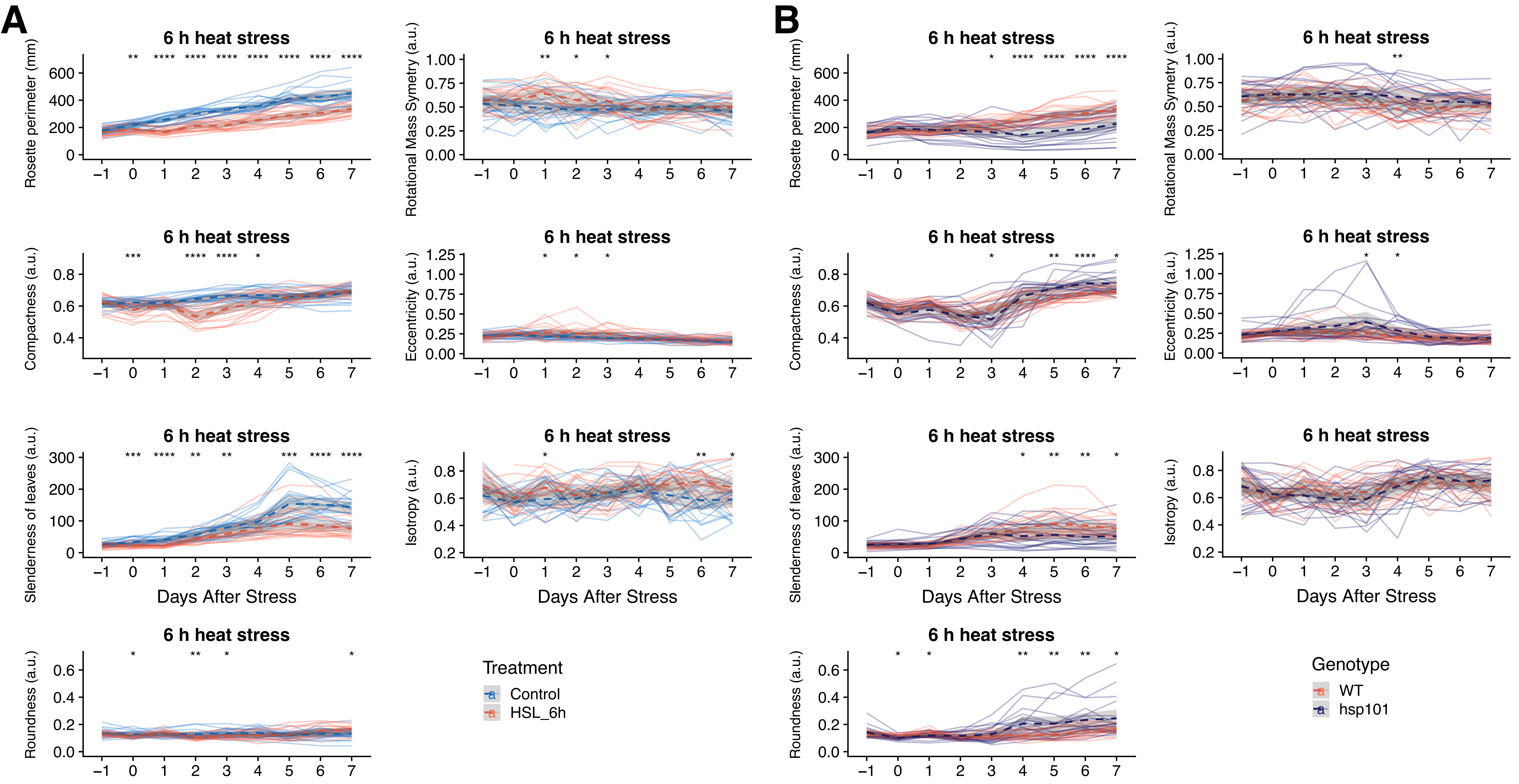


C

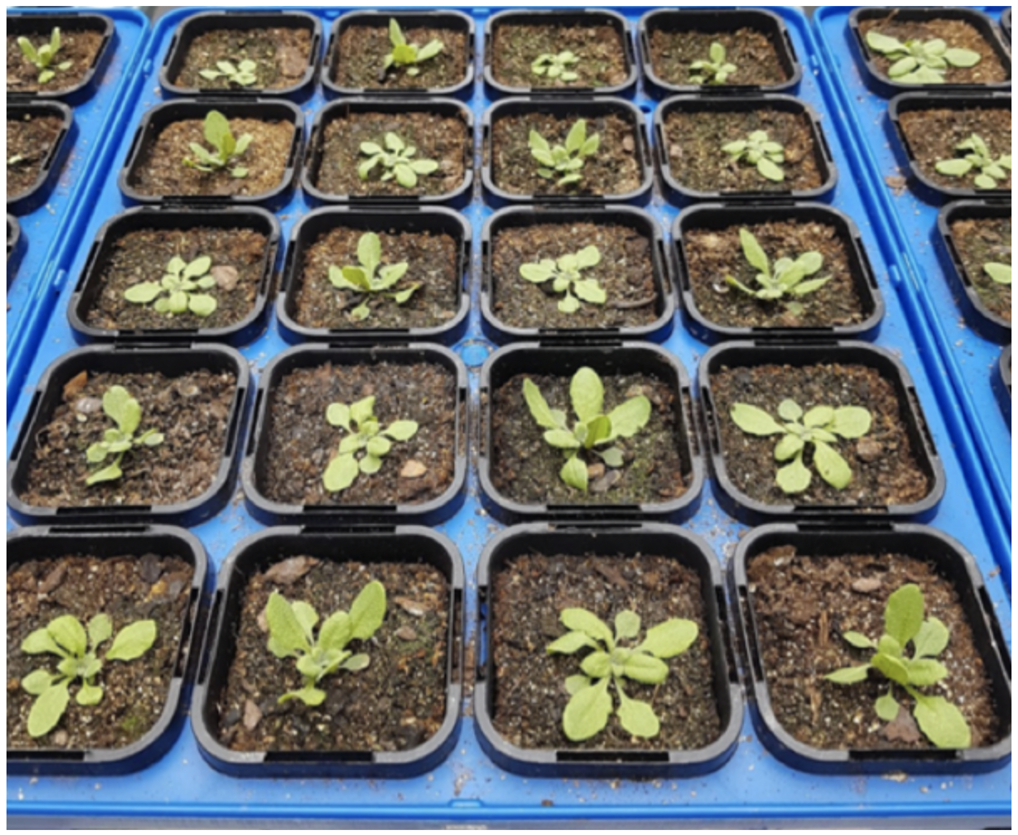

| | | | |
|---|---|---|---|
| *hsp101* | WT | *hsp101* | WT |
| WT | *hsp101* | WT | *hsp101* |
| *hsp101* | WT | *hsp101* | WT |
| WT | *hsp101* | WT | *hsp101* |
| *hsp101* | WT | *hsp101* | WT |

### Figure S3

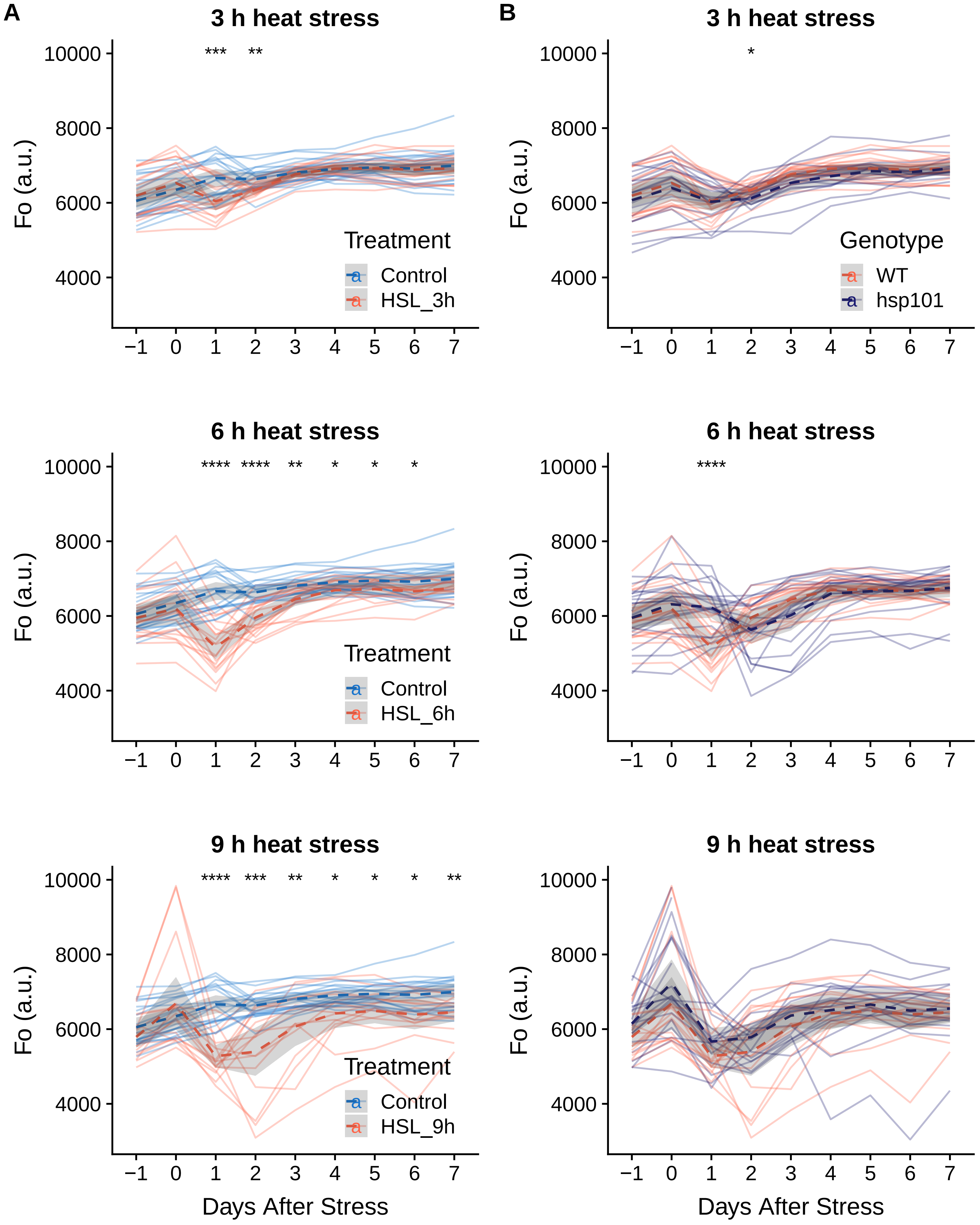

### Figure S4

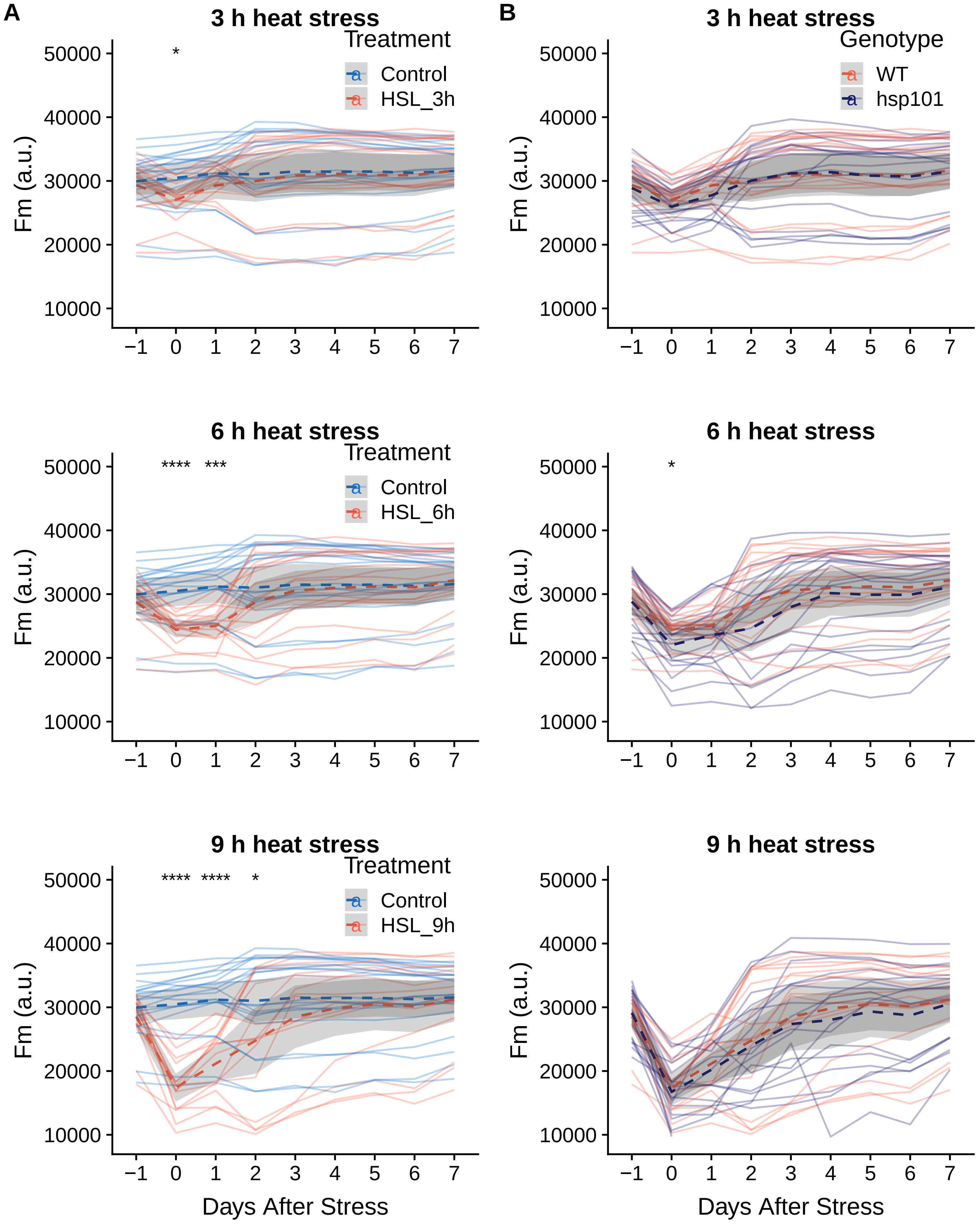

### Figure S5

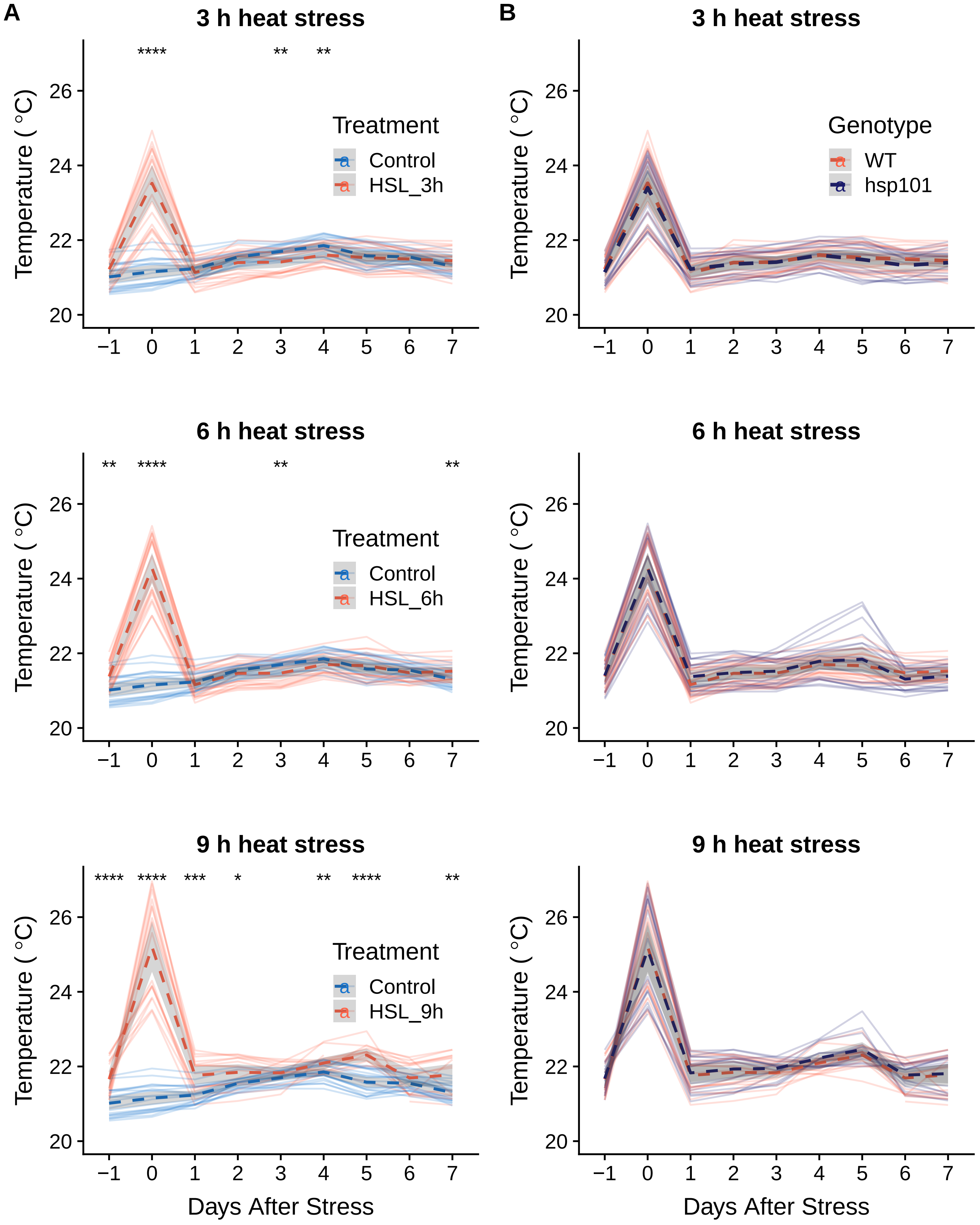

### Figure S6

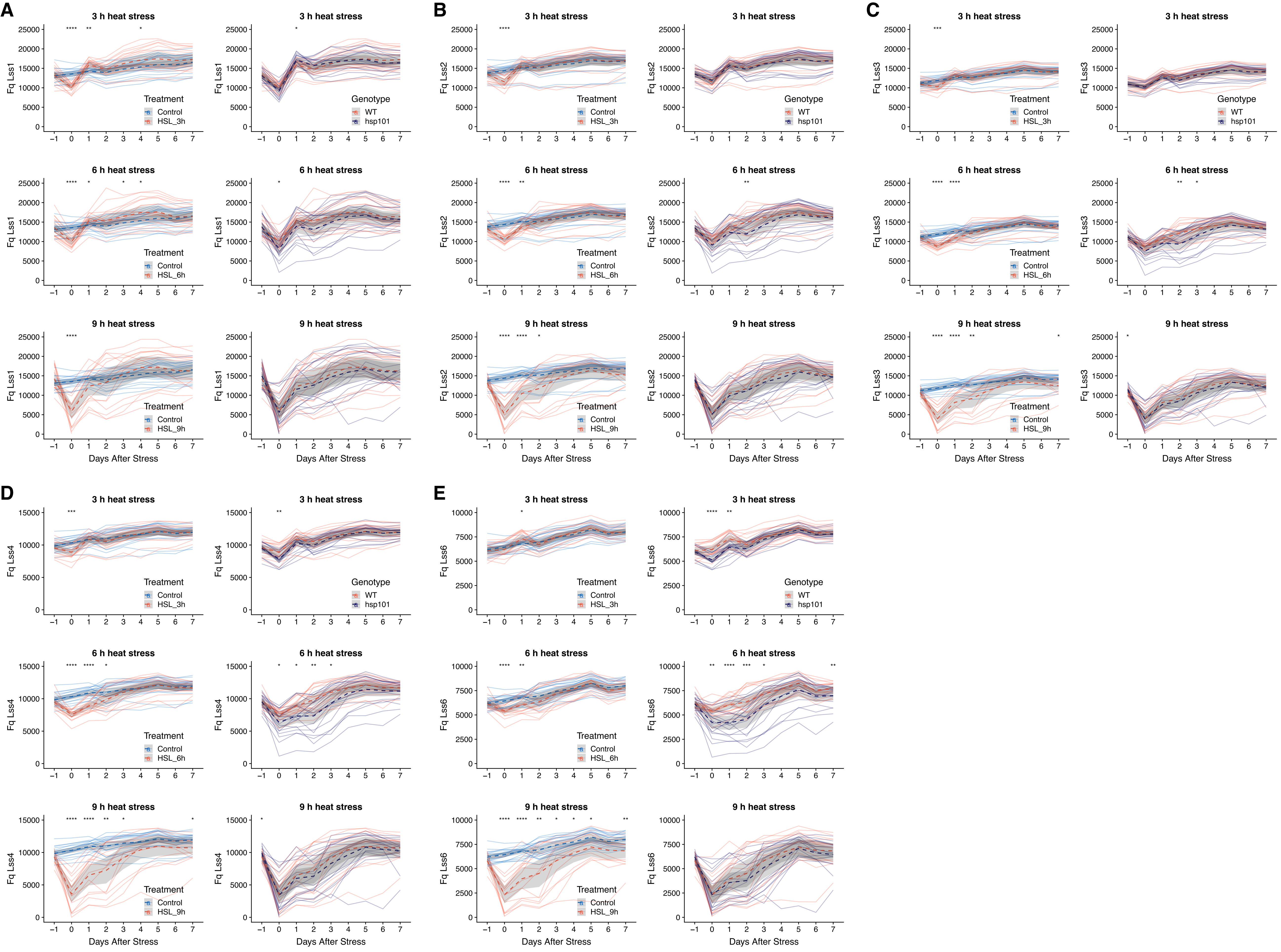

### Figure S7

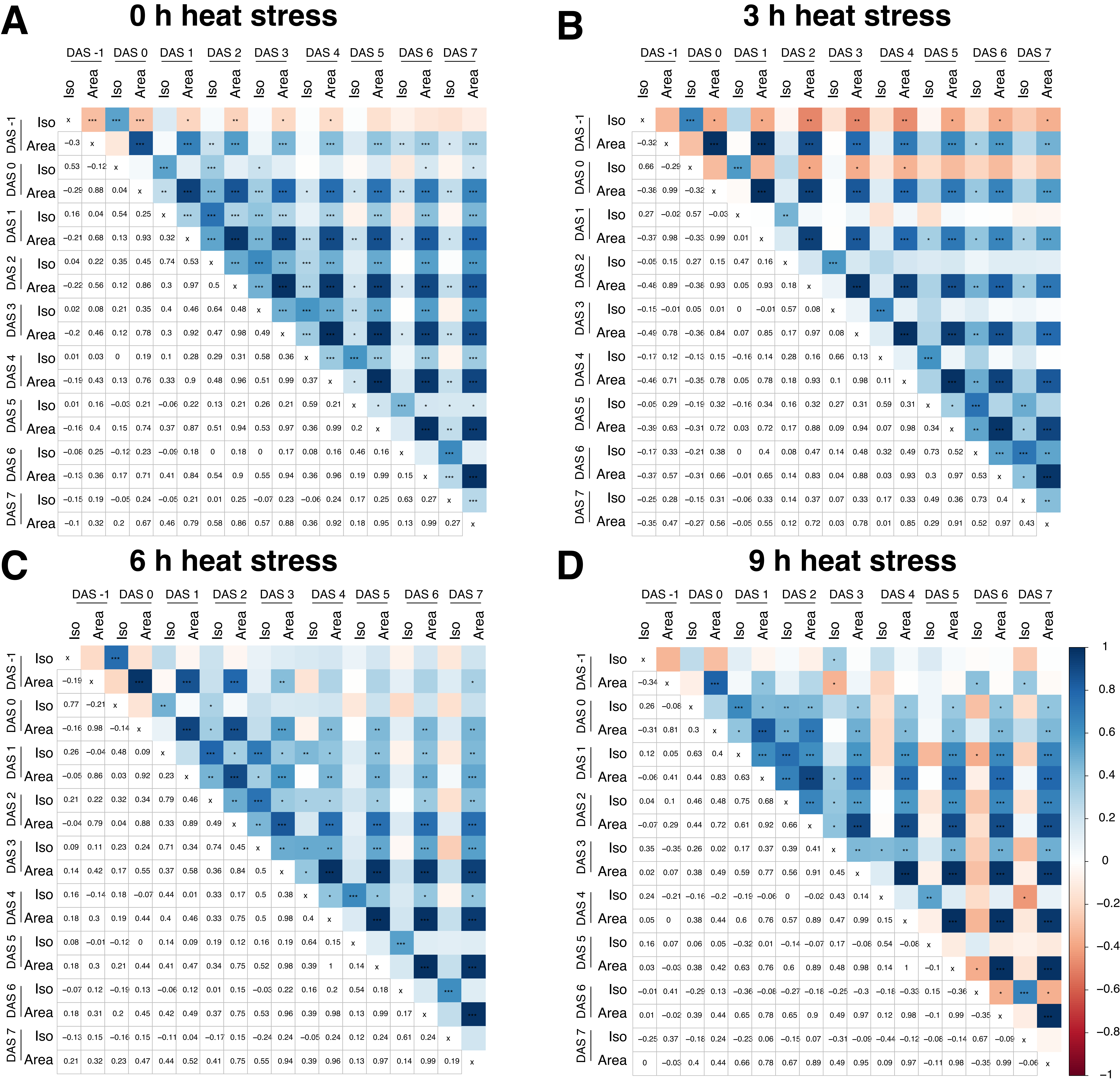

### Figure S8

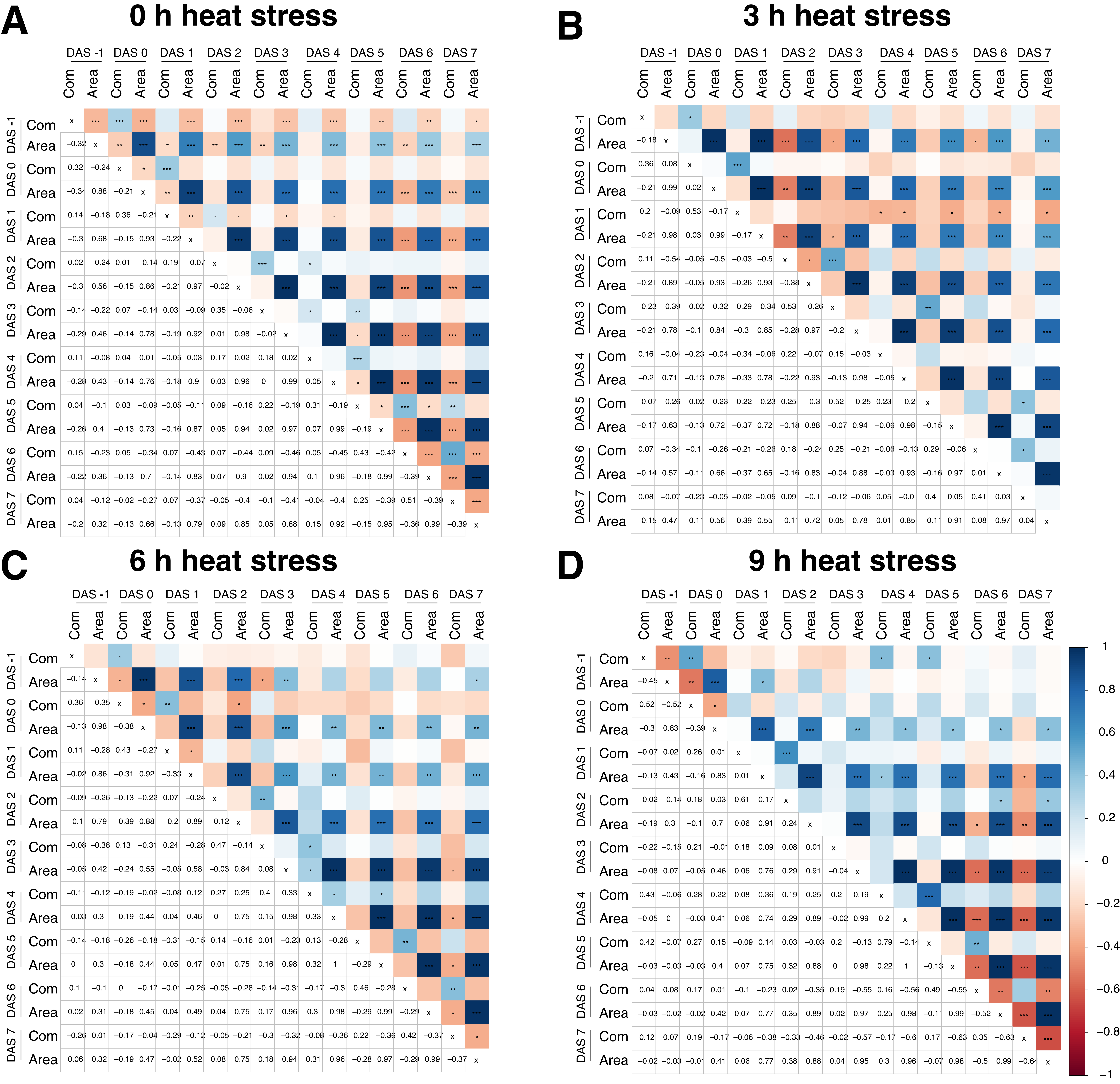

### Figure S9

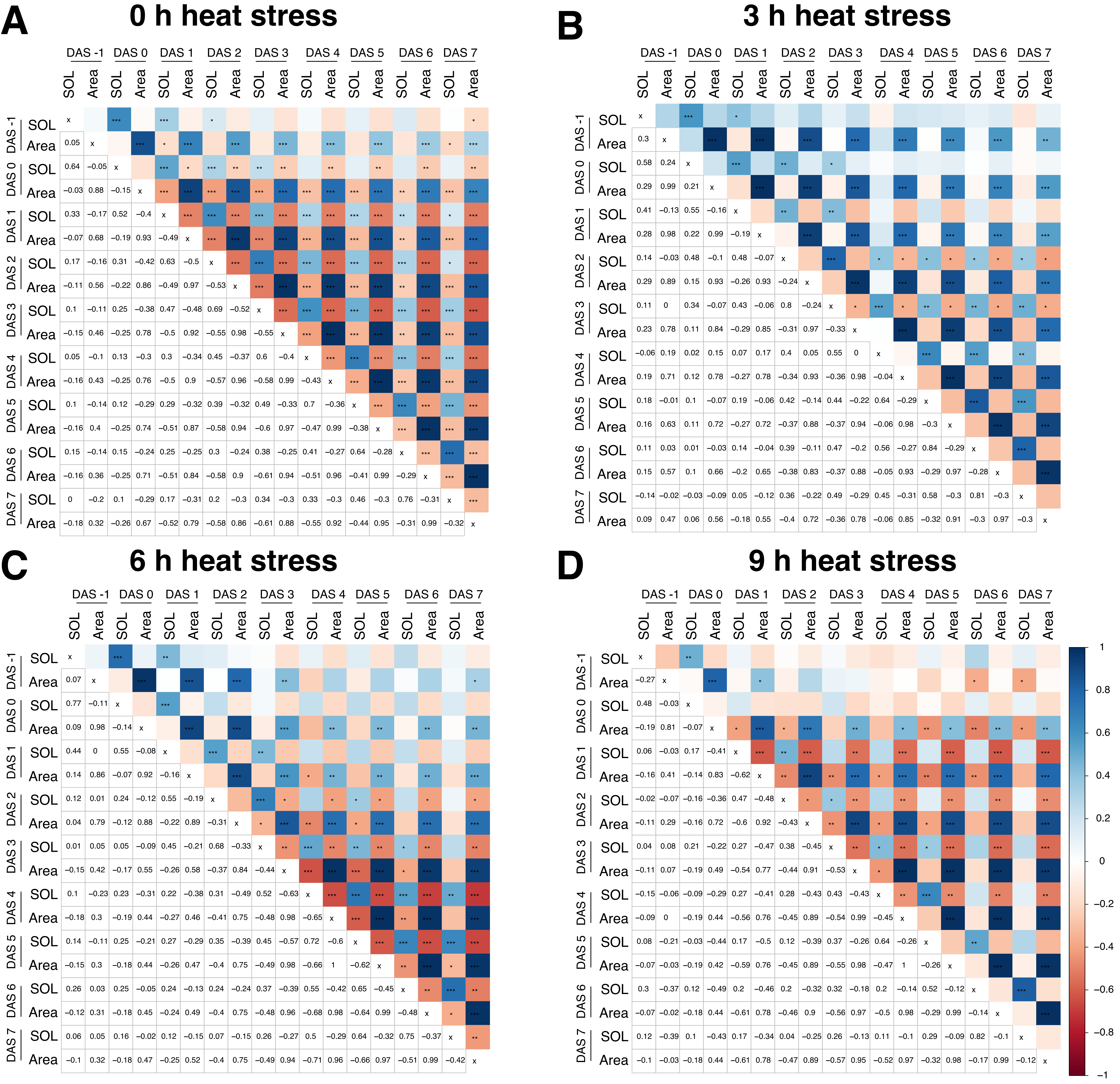

### Figure S10

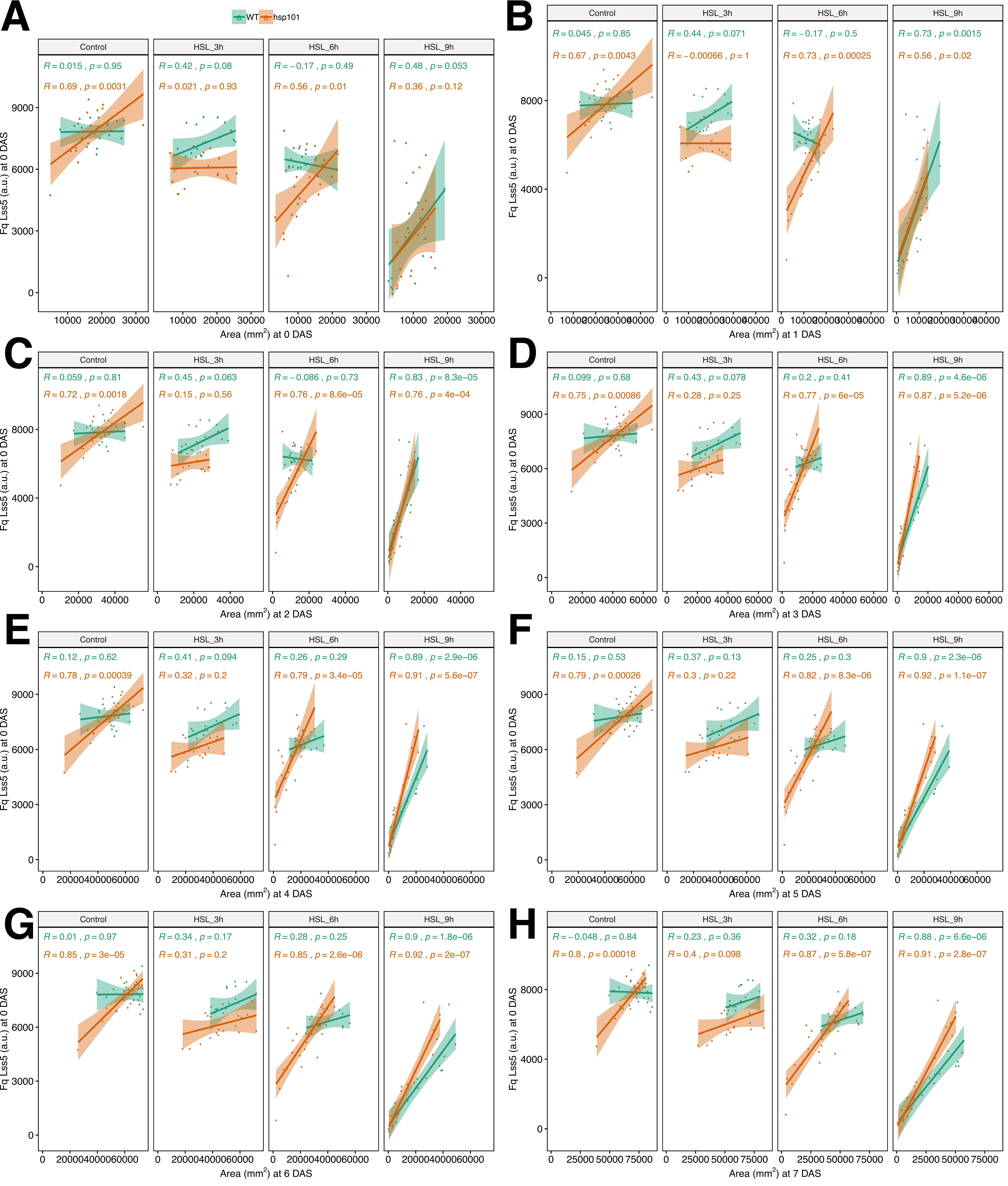

### Figure S11

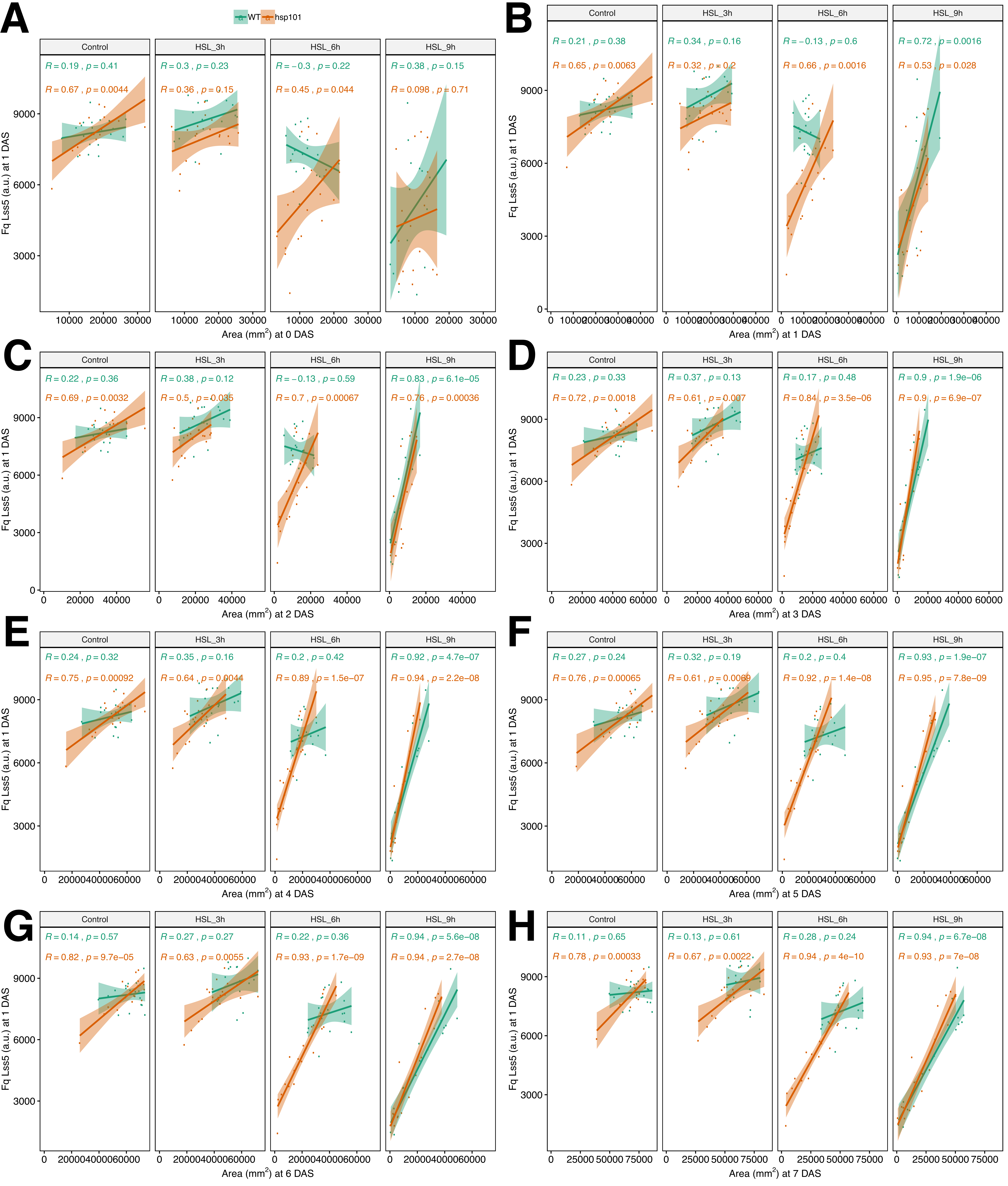

### Figure S12

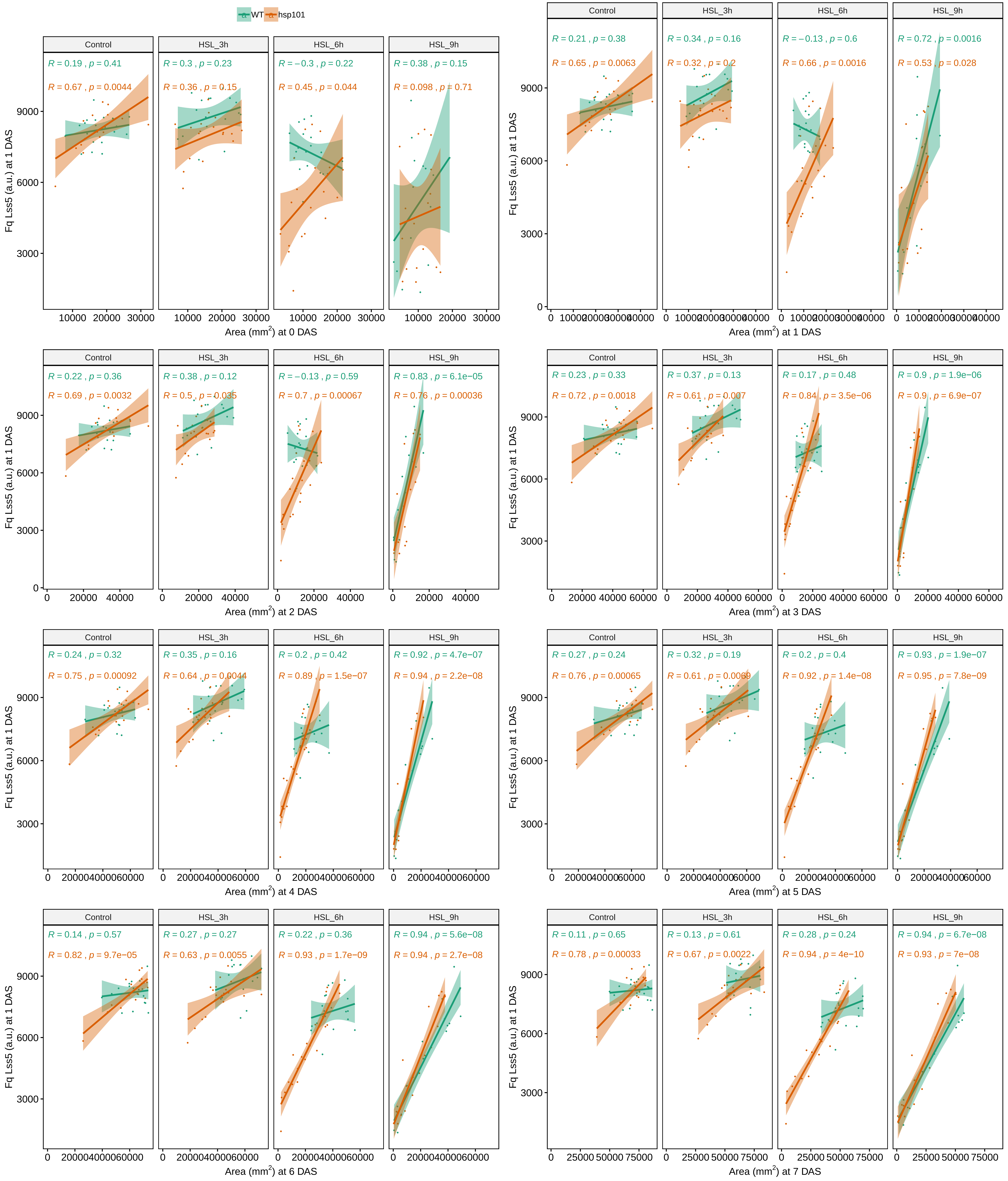
